# The exosome degrades chromatin-associated RNAs genome-wide and maintains chromatin homeostasis

**DOI:** 10.1101/2023.04.10.536209

**Authors:** Jordi Planells, Antonio Jordán-Pla, Shruti Jain, Juan José Guadalupe, Estelle Proux-Wera, Anne von Euler, Vicent Pelechano, Neus Visa

**Author notes:** Corresponding authors: Neus Visa Antonio Jordán-Pla.

## Abstract

Chromatin-associated RNAs (caRNAs) modulate chromatin organization and function. The RNA exosome degrades different types of nuclear transcripts, but its role in chromatin has not been addressed. Here we have used *Drosophila melanogaster* S2 cells as a model system to identify the repertoire of caRNAs and establish the role of the exosome in their regulation. We have analyzed both unique and repetitive sequences, and combining RNA-seq and ATAC-seq we show that the simultaneous depletion of the exosome catalytic subunits RRP6 and DIS3 not only affects caRNA levels but also changes the local chromatin accessibility at specific loci. We have identified a group of exosome-sensitive genes that are involved in developmental regulation and are characterized by a balanced chromatin state in which Polycomb and Trithorax factors coexist. Our results reveal that RNA degradation by the exosome is an important mechanism for the homeostasis of such balanced chromatin states. Given that eukaryotic genomes are repetitive to a large extent, we have also analyzed repetitive caRNAs (rep-caRNAs) and we show that the exosome is needed to control rep-caRNA levels and to maintain the degree of chromatin packaging in repetitive genomic regions. This role is particularly relevant in the pericentromeric regions where the exosome is required to silence LTR elements and maintain centromere organization.

## INTRODUCTION

Eukaryotic genomes are extensively transcribed and encode a large variety of coding and non-coding RNAs (ncRNAs). Some of these RNAs remain associated with the chromatin and participate in the regulation of fundamental biological phenomena such as gene expression, chromatin organization and DNA repair through mechanisms that are only partially understood [1,2]. Knowledge about the regulatory potential of ncRNA in the chromatin came from studies on the role of Xist in chromosome inactivation [3] and from paradoxical observations in the pericentromeric heterochromatin of *Schizzosaccharomyces pombe*, where small interfering RNAs (siRNAs) that were transcribed from silenced genomic sequences were necessary to recruit repressive histone modifying enzymes [4]. Since these early observations on the roles of ncRNAs in chromatin regulation, a plethora of other regulatory RNA species have been identified, including PIWI-interacting RNAs [5], long ncRNAs (lncRNAs) [6], enhancer RNAs (eRNAs) [7], cryptic ncRNAs that result from pervasive transcription [8,9] and DNA damage-induced RNAs (diRNAs) [10]. To what an extent these RNAs are efficiently evicted from the transcription sites or remain in the chromatin is an open question.

The RNA exosome is a multiprotein complex that degrades different types of RNA, including many ncRNAs. It consists of a nine-subunit catalytically inactive core and two catalytic subunits with ribonuclease activity that associate with the core [11]. One of the catalytic subunits, RRP6 (also known as EXOSC10) belongs to the RNase D family and has 3’-5’ exoribonuclease activity. The other one, DIS3 (also known as Rrp44 or EXOSC11) is a RNase II enzyme that has both 3’-5’ exo- and endoribonuclease activities [12]. The RNA exosome is conserved throughout evolution and is involved in many cellular processes, including pre-rRNA and pre-snoRNA processing, mRNA quality control, degradation of unstable transcripts and DNA repair [13-16]. The RNA exosome has also been reported to regulate a subset of protein coding genes in flies and mammals though mechanisms that are not fully understood [17,18].

The presence of RNA in the chromatin can influence its local properties through different mechanisms, as shown by *in vitro* and *in vivo* experiments. Chromatin-associated RNAs (caRNAs) can interact with histone tails and reduce their electrostatic interactions with the DNA, thus modulating the packaging of the nucleosome fiber [19]. CaRNAs can also bind specific non-histone proteins associated with DNA and divert them from the chromatin. Indeed, many chromatin factors have been shown to bind RNA [20,21], including a major heterochromatin component, heterochromatin protein 1 (HP1) [22], and the Polycomb repressor complex 2 [23,24]. These observations suggest that targeting ribonuclease activities to specific genomic sites for efficient RNA degradation is a necessary mechanisms of chromatin homeostasis. In support of this hypothesis, studies in *D. melanogaster* [25], *C. elegans* [26] and human cells [27] have revealed functional links between nuclear RNA degradation and chromatin control. However, the role of the exosome in regulating chromatin organization has not been investigated genome-wide. Here, we have used *D. melanogaster* S2 cells as a model system to identify the repertoire of caRNAs, understand the role of the exosome in their degradation, and establish links between caRNAs, gene regulation and chromatin compaction. We show that the exosome controls caRNA levels and chromatin accessibility in a group of developmentally regulated genes that show a “balanced” chromatin state characterized by the simultaneous presence of Polycomb and trithorax factors, which implies that caRNA degradation by the exosome contributes to fine-tune developmental genetic programs. Given that eukaryotic genomes are repetitive to a large extent, we have also analyzed repetitive caRNAs (rep-caRNAs) and our results provide evidence that the exosome is not only needed to degrade repetitive transcripts, but also to maintain the degree of chromatin packaging in repetitive genomic regions. This phenomenon is especially important in the pericentromeric regions where the exosome is required to maintain the structure of the centromeres.

## MATERIAL AND METHODS

### Culture of S2 cells

*Drosophila melanogaster* Schneider 2 (S2) cells were cultured at 28 °C in Schneider’s medium (Gibco) supplemented with 10% fetal bovine serum (Gibco), 50 μg/ml streptomycin and 50 U/ml penicillin (Gibco).

### Depletion of exosome subunits by RNA interference (RNAi)

RNAi was carried out essentially as described in [28]. The MegaScript RNAi kit was used to produce dsRNA for RNAi experiments. Primer sequences are listed in Suppl. Table S1. Cells were incubated with dsRNAs complementary to GFP (control), Rrp6, Dis3, or both (double knockdown condition) for 48 hours at 28°C. The treatment was repeated once more at 48 hours and cells were harvested 96 hours after the initial transfection. All experiments were carried out in biological triplicates.

The knock-down efficiencies were analyzed by RT-qPCR as described below. Act5C, GAPDH and 28S rRNA genes were selected as reference genes. The sequences of the primers used for qPCR analysis are presented in Suppl. Table S2.

### RNA extraction

Total RNA was extracted using Trizol (Ambion) and ethanol precipitated following standard procedures. Isolated RNA was quantified using NanoDrop One.

For chromatin-associated RNA extraction,S2 cells were crosslinked with formaldehyde for 10 min at RT and the cross-linking was stopped by addition of 1 M glycine for 10 additional min. The cells were then collected and resuspended in Buffer 1 (50 mM Hepes pH 7.6, 140 mM NaCl, 1 mM EDTA, 10% glycerol, 0.5% NP40, 0,25% Triton X100 and cOmplete protease inhibitors) and incubated for 10’ at 4 °C, followed by centrifugation at 4 °C and 500g. The pellet was resuspended in cold Buffer 2 (200 mM NaCl, 1 mM EDTA, 0,5 mM EGTA, 10 mM Tris pH 8 and cOmplete protease inhibitors) and centrifuged again. Finally, chromatin was resuspended in Buffer 3 (1 mM EDTA, 0,5 mM EGTA and 10 mM Tris pH 8 and cOmplete protease inhibitors). Chromatin was fragmented using a Bioruptor sonicator (Diagenode) with 25 high intensity sonication 30’’ on/off pulses. Protein concentration was measured using a NanoDrop ND-100 (Thermo Scientific). The chromatin extract was treated with DNAse I (Thermo Scientific) and Proteinase K (Thermo Scientific). RNA was isolated with Trizol (Ambion). RNA concentration was measured with NanoDrop.

### RT-qPCR

RNA was extracted with TRIzol reagent (Ambion, ThermoFisher), treated with 1 unit DNase I (Thermo Fisher) for 60 min and reverse-transcribed using random primers (Thermo Fisher Scientific) and SuperScript III (Invitrogen). cDNAs were used for qPCR using KAPA SYBR Fast qPCR Master Mix (Kapa Biosystems) in a QIAGEN Rotor-Gene Q. Primer design was according to MIQE guidelines. All primer pairs fulfilled quality criteria according to amplification efficiency and melting curves. The results presented are compiled data from multiple independent biological replicates, each analyzed in duplicate. For each experiment, the number of independent replicates is provided in the figure legend.

### Library preparation for Illumina RNA-seq

RNA-seq experiments consisted of biological triplicates, but the total RNA samples were pooled and sequenced together. Before RNA-sequencing, and in order to obtain adequate sequencing coverage for RNAs over such a broad size range, the RNA preparations were fractionated using AMPure XP paramagnetic beads (Agilent technologies) at a concentration of 0,6X to select RNAs larger than 500 nt and 1,6X to select caRNAs smaller than 500 nt. The 0-500 nt caRNAs were directly submitted to ribosomal depletion and library preparation, whereas caRNAs >500 nt were first fragmented by sonication. The NEBNext rRNA Depletion kit was used for ribosomal RNA depletion. Single-end strand-specific libraries were prepared using NEBNext Ultra RNA library Prep Kit for Illumina (NEB) according to the manufacturer’s instructions. Samples were sequenced in a Illumina NextSeq 500 sequencer aiming for a depth of 15 million reads in average per sample with a length of 80 nucleotides per read. The number of raw, mapped and ribosomal reads in each sample are shown in Suppl. Table S3. After sequencing, the reads for 0-500 nt and >500 nt caRNAs from each experiment were merged and analyzed together. The correlations between biological replicates were high (Spearman correlation coefficients > 0,9) (Suppl. Fig. S1).

### RNA-seq data analysis

FastQC was used for sequence quality control. Cutadapt and Trimmomatic were used pre-process raw sequencing reads. Cutadapt was used to remove Illumina adapters. The SLIDINGWINDOW function from Trimmomatic was applied to discard reads with an average quality of less than 28 and the MINLEN function was used to filter out reads shorter than 20 bases. A second round of inspection with FastQC was carried out after the filtering steps to ensure that the filters were applied properly. High quality reads were aligned against the Drosophila melanogaster DM6 genome assembly with TopHat, using default settings. Sequences from the 0-500 nt and >500 nt fractions were aligned individually and then merged with the SAMtools merge function. For visualization purposes, genome browser tracks were generated with the bamCoverage function as implemented in the deepTools suite, using the alignment files as input. The coverage values were RPKM normalized and the resulting coverage data tracks were visualized with the Integrative Genomics Viewer software (IGV). When downstream analysis did not require the samples to be considered as separate biological replicates, read alignments were merged into a single alignment dataset using SAMtools. The featureCounts function from the R package RSubread was used to extract the number of reads that mapped to the different genes. The 9-state combinatorial chromatin annotation of the genome was obtained from [29]. The R packages Voom and Edge R were used for the differential expression analysis at the gene level. Variance stabilized counts were computed with DESeq2 and used for PCA analysis. Interval operations have been performed with bedtools, GRanges and plyranges.

Normalized reads (TPM) have been computed individually for each replicate and averaged. Repetitive elements were classified according to their expression level using two different TPM thresholds. Rep-caRNAs with TPM > 0 in at least one replicate were classified as detected. Rep-caRNAs with TPM > 3 in all replicates constituted the confident rep-caRNA set. GO enrichment analysis was carried out using the function enrichGO from the R package clusterProfiler.

### Library preparation for Illumina ATAC-seq and ATAC-seq data analysis

All the samples consist of biological triplicates. Tagmentation and library preparation was performed according to [30, 31] using Nextera DNA library kit (FC-121-1030). Libraries were sequenced in NextSeq500, obtaining paired-end, unstranded 37 cycles reads. The average depth obtained was over 50.4 million mapped reads (Suppl. Table S4). The nf-core pipeline [32] version 1.2.1 was used to pre-process (alignment and quality check) the ATAC-seq reads. Briefly, fastq quality was checked with FASTQC, adapters were trimmed with TrimGalore! and reads were aligned to Dm6 genome with bwa keeping multimapping reads. Narrow peaks were called using MACS with default settings (https://github.com/nf-core/atacseq). For transcription factor binding prediction analysis, the TOBIAS pipeline [33] was used, with default options and modifying the MACS --gsize 142573017 parameter. All transcription factor DNA-binding motifs available in JASPAR2022-CORE database were used for the analysis.

### Reference genome, gene annotation and publicly available data

Reference genome and gene annotation used for the study were downloaded from ensembl database, version BDGP6.28 (Berkeley Drosophila Genome Project, [34]) and BDGP6.28.100, respectively. BDGP6.28.100 annotation was slightly modified by excluding rRNAs and tRNAs. Repetitive elements annotation was obtained from RepeatMasker Dm6 precomputed annotations (https://www.repeatmasker.org/genomicDatasets/RMGenomicDatasets.html). The following repeat classes were excluded from the analysis: RNA, rRNA, Other, SINE/tRNA-Deu-L2, Unknown. Pooled annotation used for Fig. 2A is a concatenation of both BDGP6.28.100 and RepeatMasker annotations with the aforementioned modifications. For the differential expression analysis of repetitive elements, each genomic coordinate was extended 50bp up/downstream in order to maximize the number of uniquely mapping reads. Adjacent overlapping regions were considered as one if both regions shared strand and repetitive class (Suppl. Data 1). To characterize the chromatin signatures of DE repetitive elements, ChIP-seq data for several chromatin marks and proteins were used (GEO accessions in Suppl. Table S5). Datasets aligned to the Dm3 reference genome (9-state chromatin model, modENCODE datasets) were made compatible with Dm6 alignments using CrossMap.py (0.6.3). Multiple modENCODE datasets for the same protein were merged using bigwigCompare (deeptools) by averaging the signal across bins. ChIP-seq data were analyzed with the nf-core/chipseq version (1.1.0).

### Fluorescence in situ hybridization (RNA-FISH)

Larval salivary glands were dissected and fixed with 3.6% formaldehyde in PBS containing 1% Triton X-100 for 40 sec followed by 2 min in 3.6% formaldehyde/50% acetic acid solution. The glands were squashed in lactoacetic acid (lactic acid: water: acetic acid; 1:2:3) and the coverslips flipped-off after freezing the preparations in liquid nitrogen. Chromosome squashes we prehybridized with 10 µl hybridization buffer (HB: 2xSSC, 50% formamide, 5% dextrane sulfate) for 30 min at 42° C. Probe mix was freshly prepared and contained 1.2 µl probe at 1.2 µg/ml, 0.4 µl salmon sperm DNA and 8.4 µl HB. Hybridization was for 1 h at 42° C. Slides were washed sequentially with the following solutions, 3 min each: 50% formamide at 42° C, 5 x SSC at 42° C, 2 x SSC at 42° C, 0.1 x SSC at room temperature, and 1 x PBS/10% Tween 20 (PBT) at room temperature. The slides were then blocked with 3% BSA in PBT for 45 min and incubated with primary antibody (2 µg/ml sheep-anti-DIG in PBT). After washing in PBT twice for three minutes each, slides were incubated with secondary antibody (10 µg/ml donkey-anti-sheep-Alexa555) for 40 min at 37° C. Slides were rinsed again with PBT and mounted with 10 µl Vectashield (with 50% DAPI). The preparations were counterstained with an antibody against H3K9ac (Abcam ab10812). The slides were examined in a Axioplan fluorescence microscope (Carl Zeiss). The probe sequences were: J5 (Simple Repeat) CAC-ACA-CGT-AAA-CCT-AAC-ACA-CGC-ACA-GAC-GC; J7 (LINE) TAC-AGA-AAT-GAC-AGG-AAG-GAA-AGT-AGG-GGA-GG; J8 (LTR) TGC-GAC-AAC-ATA-CAA-TTC-CTG-AAA-GAT-AGT-AA.

### CID immunoflourescence

S2 cells were adhered to poly-L-lysine coated glass slides and fixed in 3,7% formaldehyde in PBS for 10 min, permeabilized in 0,1% Triton X-100/PBS for 15 minutes and blocked in a solution containing 5% milk, 5% BSA in PBS for 1 hour. An anti-CID antibody (ab10887) and a secondary antibody conjugated to Alexa Fluor 594 were used according to standard procedures. Coverslips were mounted with Vectashield containing DAPI and the slides were examined in Axioplan fluorescence microscope (Carl Zeiss).

### Immunoprecipitation and LC/MS-MS

Immunoprecipitation and high-performance liquid chromatography/tandem mass spectrometry (LC/MS-MS) were performed by Hessle et al. 2009 [35] using nuclear protein extracts prepared from S2 cells that expressed V5-tagged RRP6, SKI6 and RRP4, or from “empty” control cells that were cultivated under the same conditions but did not express any V5-tagged protein. Three independent immunoprecipitation experiments were carried out, each including an immunoprecipitation with “empty” cells processed in parallel. An enrichment probability value compared to negative controls was calculated for each protein identified using a standard error model and protein abundances were normalized by the abundance of the immunoglobulin heavy-chain variable region [*Mus musculus*] (gi 27728890). Significant interactions were defined by a false discovery rate (FDR) threshold of 0.01 and a minimum fold-difference of 2 compared to “empty” cells. The list of RRP6 interactors was provided by Eberle et al. 2015 [25]. SKI6 and RRP4 interactors are provided in Suppl. Table S6.

### Software

All tools used in the present study are publicly available. For a complete list of tools and versions used, see the Suppl. Table S7.

### Statistical analysis

Histograms show average values and error bars represent standard deviations. The numbers of biological replicates are indicated in the figure legends. T-tests were used for statistical testing of results from single gene analyses. The Kolmogorov-Smirnov (K-S) test was used for statistical testing of differences between metagene distributions. Statistical testing of GO enrichments was carried using the function enrichGO from R package clusterProfiler. The tests used in each case are indicated in the figure legends. All plots and statistical analysis have been performed with the indicated tools (Suppl. Table S7) and R software (4.2.2).

### Data availability

The RNA-seq and ATAC-seq data produced in this study are available from NCBI Gene Expression Omnibus (GEO accession GSE222262). Accession numbers to other publicly available datasets used here are provided in Suppl. Table S5.

## RESULTS

### The chromatin has a specific RNA composition different from that of total RNA

To assess the role of the exosome in the regulation of caRNAs, we used RNA interference (RNAi) to deplete exosome subunits in S2 cells and characterized the chromatin-associated transcriptome of control and exosome-depleted cells. The two exosome catalytic subunits were knocked down either individually (RRP6-kd or DIS3-kd) or simultaneously (Double-kd), and we used a dsRNA complementary to GFP as a control in the RNAi experiments. The extent of the depletions is shown in Suppl. Fig. S2. After the RNAi treatment, the cells were fixed with formaldehyde, chromatin was isolated and RNA was extracted from the chromatin fractions. Total RNA was also extracted and analyzed in parallel for reference purposes. Chromatin enrichment of a specific RNA known to be associated with chromatin was assessed by RT-qPCR (Suppl. Fig. S3). The RNA extracted from chromatin was shown to contain transcripts encompassing a wide range of lengths from 30 to 3000 nt (Fig. 1A). CaRNAs from three independent RNAi experiments were sequenced.

**Figure 1.**
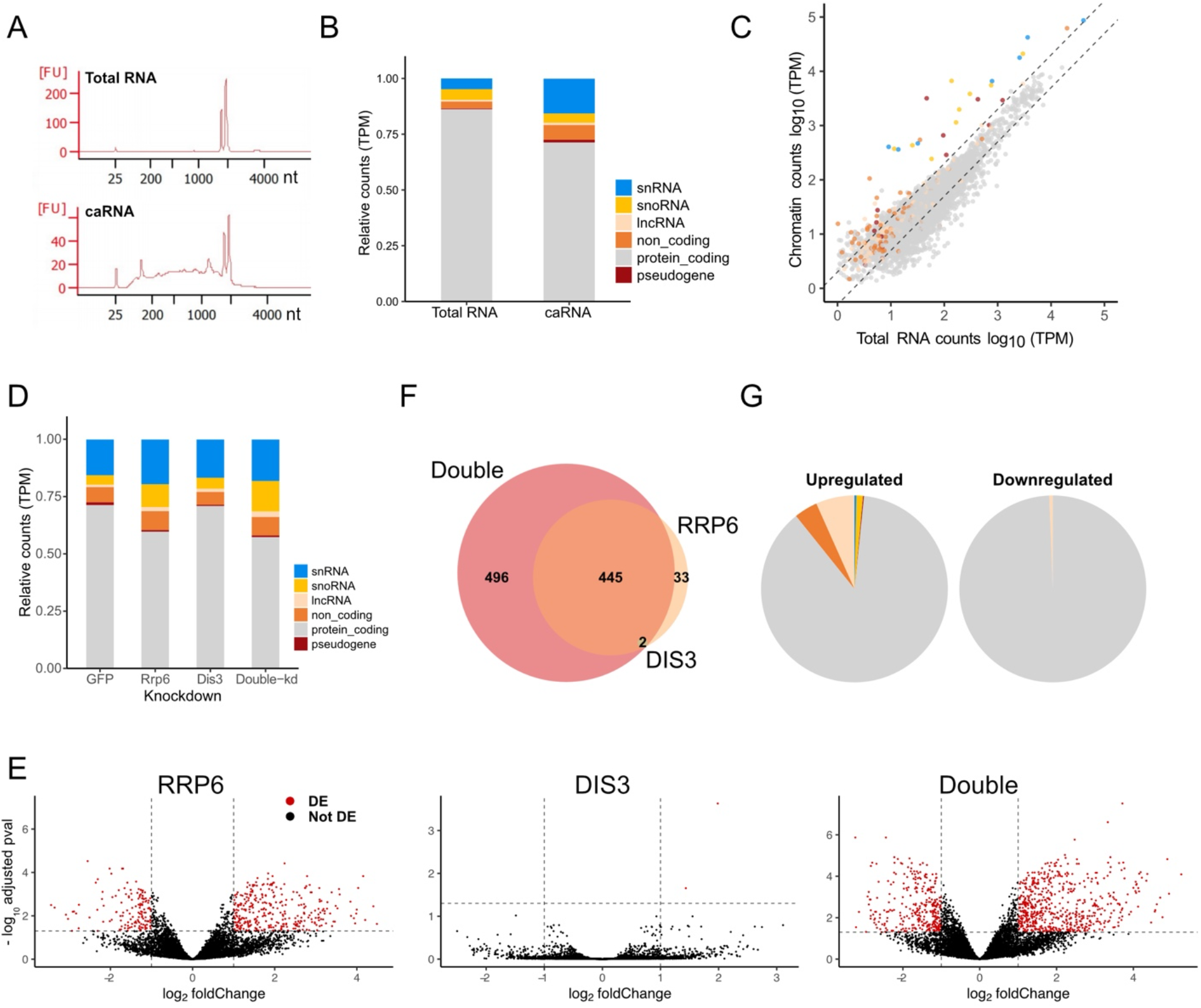
Sequencing of chromatin-associated RNAs and effects of exosome depletion on the chromatin-associated transcriptome. **(A)** Capilary electrophoresis profiles of RNAs purified from total RNA preparation (top) and chromatin fraction (bottom). **(B)** The bar plots show the relative distribution of normalized transcript abundances (TPM) per transcript biotype corresponding to unique sequences in total RNA and chromatin RNA fractions in control conditions, as indicated. The y axis shows the distribution of normalized counts (TPM). In chromatin fraction, reads were averaged across replicates. **(C)** Comparison of RNA abundance in total and chromatin preparations.The scatter plot shows log10 transformed normalized counts in total RNA preparations (x axis) versus log10 transformed normalized counts in chromatin fractions (y axis). The colors represent the different transcript biotypes as in B. Transcripts with no measurable expression in any of the conditions were removed from the analysis. The dotted black lines demarcate the +/- 2 fold interval. **(D)** RNA exosome catalytic subunits were depleted by RNAi, as indicated in the x-axis. The bars show the distribution of stacked TPM normalized counts per gene biotype in the different conditions of the study. Read abundances were normalized for each sample individually and then averaged through replicates. Control corresponds to dsGFP-treated cells. **(E)** Differential expression analysis of individual RNA exosome subunits depletion (RRP6, left panel; DIS3, middle panel) and both RNA exosome subunits (Double-kd, right panel) versus control S2 cells. The volcano plots show the average log_2_ fold change (x-axes) for each unique transcript. The y axes show the inverse of log_10_ p-value adjusted. The vertical dotted lines denote log_2_FC = -1, 1. The horizontal dashed lines correspond to the signaficance threshold used for differential expression analysis, -log_10_(0.05). Differentially expressed genes are marked with red dots (adjusted p-value <0.05 and log2FC > Ι1Ι. Total number of genes included in the analysis n=10319. **(F)** Venn diagram showing the intersection of significant DE caRNA genes detected in each of the three depletion conditions tested versus control. **(G)** Pie charts showing the gene biotypes of the DE caRNAs in Double-kd cells. Left and right plots show genes with increased and decreased RNA abundance, respectively. Colors denote biotypes as indicated in panel A.

We first analyzed the control samples (GFP control) to describe the chromatin-associated transcriptome under control conditions. The most predominant type of transcripts found in the chromatin was protein-coding RNAs, presumably corresponding to the nascent pre-mRNAs. SnRNAs, snoRNAs and other ncRNAs with known functions in the cell nucleus were also relatively abundant in the chromatin (Fig. 1B).

The relative RNA abundances in chromatin were in general well correlated with the abundances in total RNA (Spearman correlation coefficient = 0.873) although a large fraction of transcripts, mainly protein-coding, were less abundant in the chromatin fraction than in the total RNA (Fig. 1C). Altogether, these results provided a description of the specific RNA composition of the chromatin and showed that transcript abundances in chromatin and total RNA are different from each other.

### The exosome is required to maintain caRNA levels and chromatin accessibility

The analysis of caRNAs from exosome depleted cells revealed that both RRP6-kd and Double-kd had a stronger impact on the chromatin-associated transcriptome than DIS3-kd as shown by correlation and principal component analyses (Suppl. Fig. S4A and S5A). Depletion of RRP6 resulted in an accumulation of different types of non-coding RNA sequences in chromatin (Fig. 1D). A differential expression analysis versus GFP-control identified 480 and 943 differentially expressed genes (DEGs) in RRP6-kd and Double-kd, respectively, while only two genes changed in DIS3-kd (Fig. 1E-F and Table 1). Most DEGs (92%) were protein-coding genes (Fig. 1G).

**Table 1.** Number of caRNAs differentially expressed in exosome-depleted cells.

| Knockdown | Annotation | Increased | Decreased |
| --- | --- | --- | --- |
| RRP6 | Unique | 334 | 146 |
| DIS3 | Unique | 2 | 0 |
| Double | Unique | 613 | 330 |
| RRP6 | Repetitive | 204 | 20 |
| DIS3 | Repetitive | 2 | 0 |
| Double | Repetitive | 544 | 112 |

Previous studies suggested that RNA can promote chromatin decondensation [19,25,26] and we wondered whether the changes in caRNA levels observed in exosome-depleted cells would be accompanied by changes in chromatin accessibility. To investigate this possibility, we performed ATAC-seq in the same four conditions of the RNA-seq experiment (Suppl. Fig. S6 and S7, Suppl. Table S4). The single knockdowns had minor effects on chromatin accessibility, but the Double-kd had striking effects. We found 5105 differentially accessible regions (DARs) between the Double-kd and control samples. Of those, 80 % (4018 out of 5105) displayed increased accessibility compared to control (Fig. 2A). The annotation of DARs to genomic features revealed that promoters, introns and exons were predominant while only 20% of the DARs were located in intergenic regions (Fig. 2B). This observation is in agreement with previously published immunofluorescence and ChIP-seq experiments showing a strong association of the exosome with transcriptionally active loci [25,36,37].

**Figure 2.**
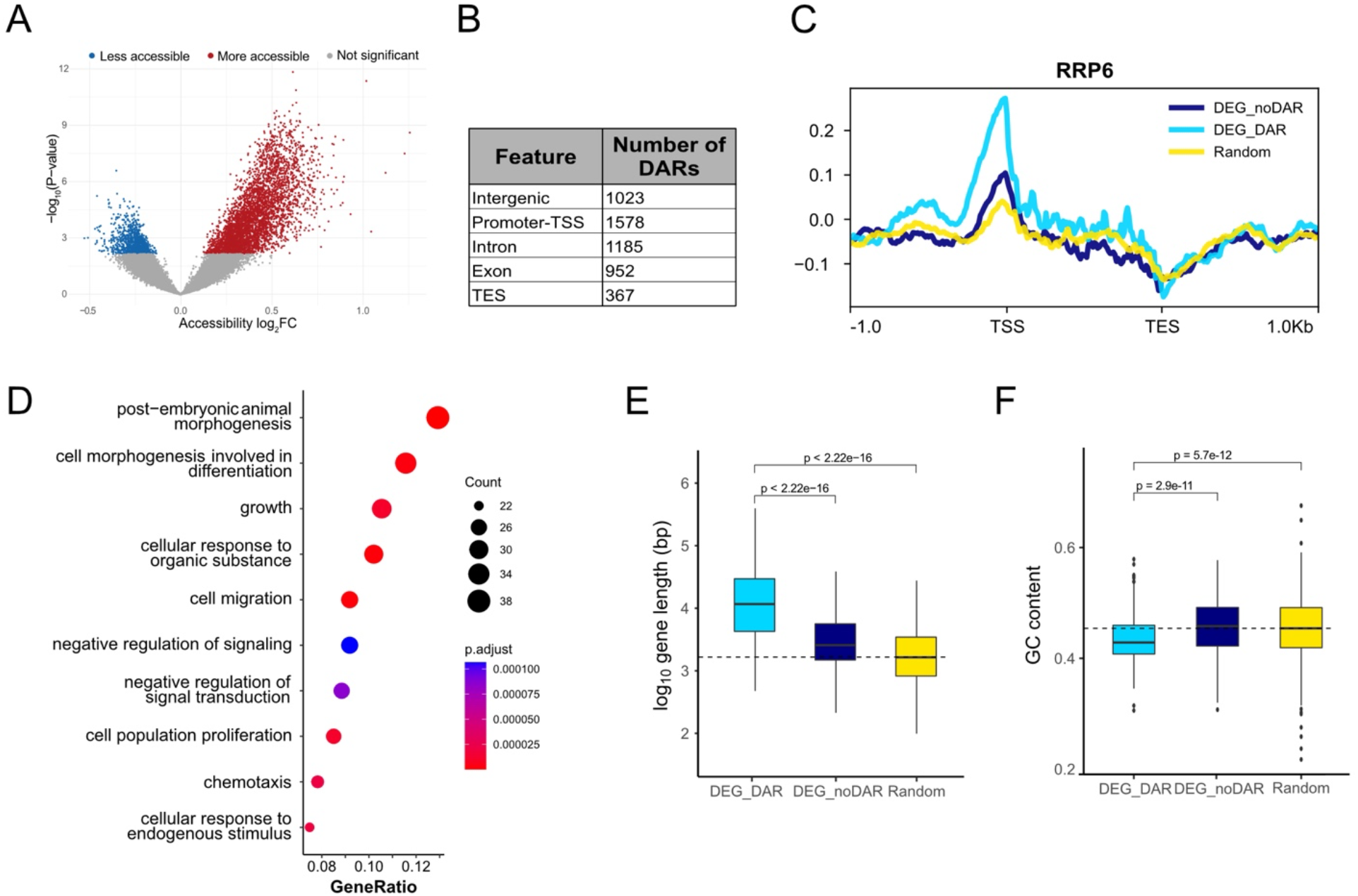
Chromatin accessibility changes induced by exosome depletion. **(A)** Volcano plot showing the result of the differential accessibility analysis in Double-kd versus control cells. The x-axis represents the average chromatin compaction change (log_2_ fold change), including data from three knockdown replicates. The y-axis represents the inverted adjusted p-value of the differential expression analysis, log-transformed. Differentially accessible regions (DARs) are indicated as blue and red dots (adjusted p-value <0.05). **(B)** Association of the DARs shown in (A) with gene features. TSS and TES denote transcription start and termination sites, respectively. **(C)** ChIP-seq profile of RRP6 average signal across three different gene sets. Light blue profile corresponds to DE caRNA genes in Double-kd intersecting a DAR (DEG_DARs, n = 352). Dark blue profile corresponds to DE caRNA genes not intersecting DARs (DEG_noDARs, n = 591). Yellow profile corresponds to a random control set of not differentially expressed caRNA genes (n=943, the sum of DEG_DAR and DEG_noDAR genes). Reads were normalized with sequencing depth. Kolmogorov-Smirnov test for DEG_DAR vs. DEG_noDAR, KS=2.95*10^-13^. **(D)** GO-analysis of molecular functions of DEG_DARs. The plot shown the top ten most significant categories in the y-axis. Statistical significance is color coded. The x-axis shows the relative amount of DE caRNA genes in each category. Dot size indicates the number of genes in each GO term. **(E)** The box plot shows the distribution of gene length for the same three sets of genes. Black line inside the boxes denotes the median length of each gene group. Two-sided non-parametric Wilcox test was used to compare the gene sets. **(F)** The box plot shows the distribution of GC content for the three gene sets. Black line inside the boxes denotes the median length of each gene group. Two-sided non-parametric Wilcox test was used to compare the gene sets.

Next, we wanted to establish whether the changes induced by exosome depletion in the local chromatin compaction were related to RNA accumulation. We intersected the genomic coordinates of DEGs and DARs and found that almost 40% of the DEGs (352 out of 943) intersected with DARs. This group of DEGs with differentially accessible chromatin in exosome depleted cells will hereon be referred to as “DEG_DARs” (see examples in Suppl. Fig. S8). The remaining 60% of DEGs were referred to as “DEG_noDARs” and did not show significantly changed chromatin accessibility. This result suggested a complex relationship between RNA levels and chromatin accessibility and showed that an increase in caRNA levels was not *per se* a determinant of chromatin accessibility. In order to understand the relationships between caRNA levels and chromatin accessibility in the two groups of genes DEG_DARs and DEG_noDARs, we measured RRP6 occupancy in previously published RRP6 ChIP-seq data [25]. DEG_DARs had a much higher RRP6 signal than DEG_noDARs (Fig. 2C, p-value =2.95*10e-13), which suggests that changes in caRNA levels at DEG_DAR loci are direct. In summary, the findings reported here show that the exosome is preferentially associated with a subset of gene loci where it regulates caRNA levels and contributes to maintain the local packaging of the chromatin.

### The exosome is a component of a complex chromatin regulatory network together with Polycomb and trithorax proteins

We further characterized the DEG_DARs to better understand the function of the exosome in their regulation. A gene ontology (GO) analysis showed that DEG_DARs are linked to morphogenesis and development (Fig. 2D). As many as 72% of DEG_DARs showed higher caRNA levels in Double-kd that in control cells, and a GO analysis for the upregulated DEG_DARs identified biological processes such as morphogenesis and development, as for the total of DEG_DARs. The downregulated DEG_DARs were instead linked to cell cycle and metabolism (Suppl. Fig. S9).

The level of expression of DEG_DARs in control conditions was not significantly different from that of DEG_noDARs (p=0.39) but, at the structural level, DEG_DARs were significantly longer and more AT-rich than DEG_noDARs (Fig. 2E-F).

A search for sequence motifs enriched in DARs identified the binding sites for the dosage compensation factor Clamp, the Trithorax transcription factor Trl/GAF and the boundary element associated factor BEAF-32 (Suppl. Fig. S10). In agreement with this observation, and using publicly available ChIP-seq data for these three factors, we observed a remarkable enrichment of Clamp and Trl/GAF in the promoter region of the DEG_DARs that was significantly higher than that observed in DEG_noDARs or in a subset of randomly selected genes (Fig. 3A), which revealed that DEG_DARs have a specific chromatin configuration.

**Figure 3.**
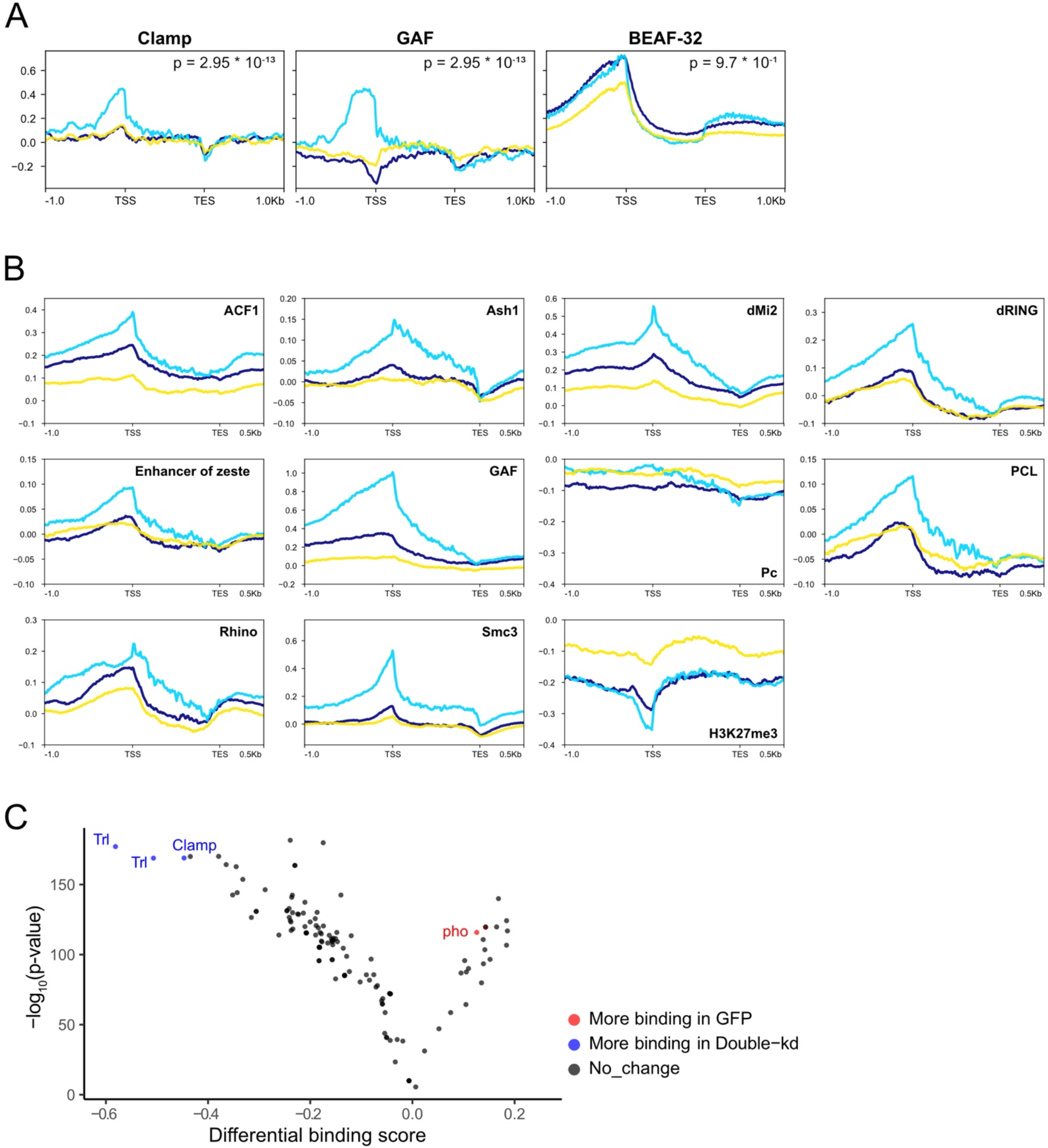
Characterization of the DEG_DAR chromatin environment. **(A)** ChIP-seq profiles of Clamp, Trl/GAF and BEAF-32 average signal across previously defined gene sets (DEG_DAR, DEG_noDAR and random set). Reads have been normalized with sequencing depth. **(B)** Chromatin binding profiles of chromatin factors with differential occupancy in DEG_DARs and DEG_noDARs. Except for H3K27me3, the plots show ChIP-on-chip data from modENCODE. The selected proteins showed statistically significant higher occupancy in the TSS region (+/- 500bp) of the DEG_DARs than in the DEG_noDARs (Kolmogorov-smirnov test, p value < 0.0001). H3K27me3 data is ChIP-seq from GSE41440 and GSE93100 (not significant). **(C)** Transcription factor binding prediction from accessibility data analysis (see Material and Methods). The volcano plot shows the TOBIAS differential binding score (x-axis) and significance as -log_10_ p-value (y-axis). Each dot represents the binding activity of the TFs obtained from JASPAR database. Top 10 most bound TF for Double-kd and control condition are colored in blue and red respectively.

We analyzed ChIP-on-chip data available at modENCODE for 40 non-histone chromatin factors (Suppl. Fig. S11) to further define the DEG_DAR chromatin landscape, and we identified ten proteins, including Trl/GAF, that were significantly more enriched in DEG_DARs than in DEG_noDARs (Fig. 3B). These proteins were all related to transcription regulation, chromatin remodeling or histone modifications, and four of them (Pc, PCL, dRING, Enhancer of zeste) were members of the Polycomb group, which was unexpected given that DEG_DARs are active genes in S2 cells. Another enriched protein at DEG_DARs was Smc3, a subunit of the cohesin complex that has been shown to mediate the recruitment of Polycomb to active genes [38]. Interestingly, the Trithorax group proteins Trl/GAF and Ash1, known for their roles in counteracting Polycomb silencing, were also significantly enriched at DEG_DARs (Fig. 3B). Moreover, analysis of H3K27me3 ChIP-seq data at DEG_DAR loci revealed low levels of H3K27me3 (Fig. 3B), in agreement with DEG_DAR loci not being repressed by Polycomb. These findings suggest that DEG_DAR loci are regulated by a complex interplay between Polycomb and Trithorax proteins, and that the exosome is a component of this regulatory network. This conclusion was further supported by the result of a differential transcription factor DNA-binding prediction analysis based on the ATAC-seq signals (TOBIAS, see Material and Methods for details) (Fig. 3C). This analysis identified 143 genes that were bound by both Trl/GAF and Pleiohomeotic (Pho), a DNA-binding component of the *Drosophila* Polycomb complex [39], in control S2 cells (Suppl. Table S8). TOBIAS also predicted increased Trl/GAF chromatin binding and decreased binding of Pho in Double-kd. This change in the overall binding of Trithorax and Polycomb factors in response to RRP6 and DIS3 depletion links the exosome to the regulation of balanced chromatin states.

Next, we searched for protein-protein interactions that could provide support for a physical link between the exosome, Polycomb and Trithorax factors. In previous studies, we stably transfected S2 cells with plasmids that expressed V5-tagged exosome subunits and carried out coimmunoprecipitation experiments to identify exosome-binding partners by high-throughput mass spectrometry [25,35]. By further mining the interaction network of the exosome we identified interactions between RRP6 and Su(z)12, a component of the PCR2, and between the exosome core subunit SKI6 and both dRING and Trl/GAF (Suppl. Table S6).

In summary, we have shown that the exosome is required together with Polycomb and Trithorax proteins to regulate the chromatin state at specific gene loci (DEG_DAR loci) in the *Drosophila* genome. These genes are transcriptionally active in S2 cells and are involved in important developmental processes such as morphogenesis, cell differentiation and proliferation. The coexistence of Polycomb with Trithorax factors and active marks in transcriptionally active genes has been previously reported and proposed to constitute a ‘balanced’ chromatin state that can be modulated by both repressors and activators during development [40]. Our results identify RNA degradation by the exosome as an important mechanism for the homeostasis of such balanced chromatin states.

### Repetitive RNAs are also components of the chromatin-associated transcriptome

Next, we aimed at characterizing the repetitive chromatin-associated transcriptome and understanding the role of the exosome in its regulation. We first studied the repetitive caRNAs (rep-caRNAs) in control S2 cells. If only uniquely mapped reads were included in the analysis, as in conventional RNA-seq pipelines, a total of 10 872 expressed rep-caRNAs were detected, which represented only 11,3 % of the chromatin-associated transcriptome (Fig. 4A, left). In a more comprehensive quantification of repeat abundance including the multimapping reads that aligned to the RepeatMasker annotation (see Material and Methods), the number of detected rep-caRNAs in control cells was almost twice as high, 19 458 (Table 2 and Fig. 4A, middle). This strategy did not assign each read to its locus of origin, but provided information about the types and abundances of rep-caRNAs (Fig. 4A). In total RNA preparations, repetitive transcripts were more abundant than in the chromatin and accounted for almost 50% of the transcriptome of S2 cells (Fig. 4A, middle and Table 2), which implies that rep-caRNAs are not particularly enriched in the chromatin. When only rep-caRNAs derived from intergenic sequences were considered (Fig. 4A, right), rep-caRNAs constituted only 11.3 % and 7.8% of the total and chromatin-associated transcriptomes, respectively, which indicated that most rep-caRNAs were produced from annotated gene loci.

**Table 2.**
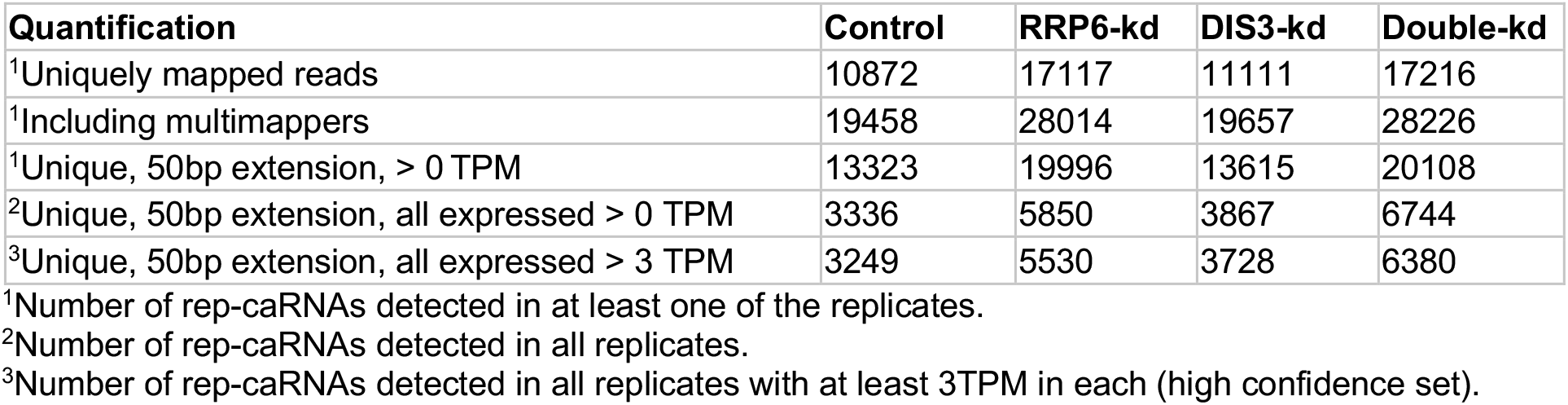
Number of rep-caRNAs detected in control and exosome-depleted cells using different RNA quantification approaches.

**Figure 4.**
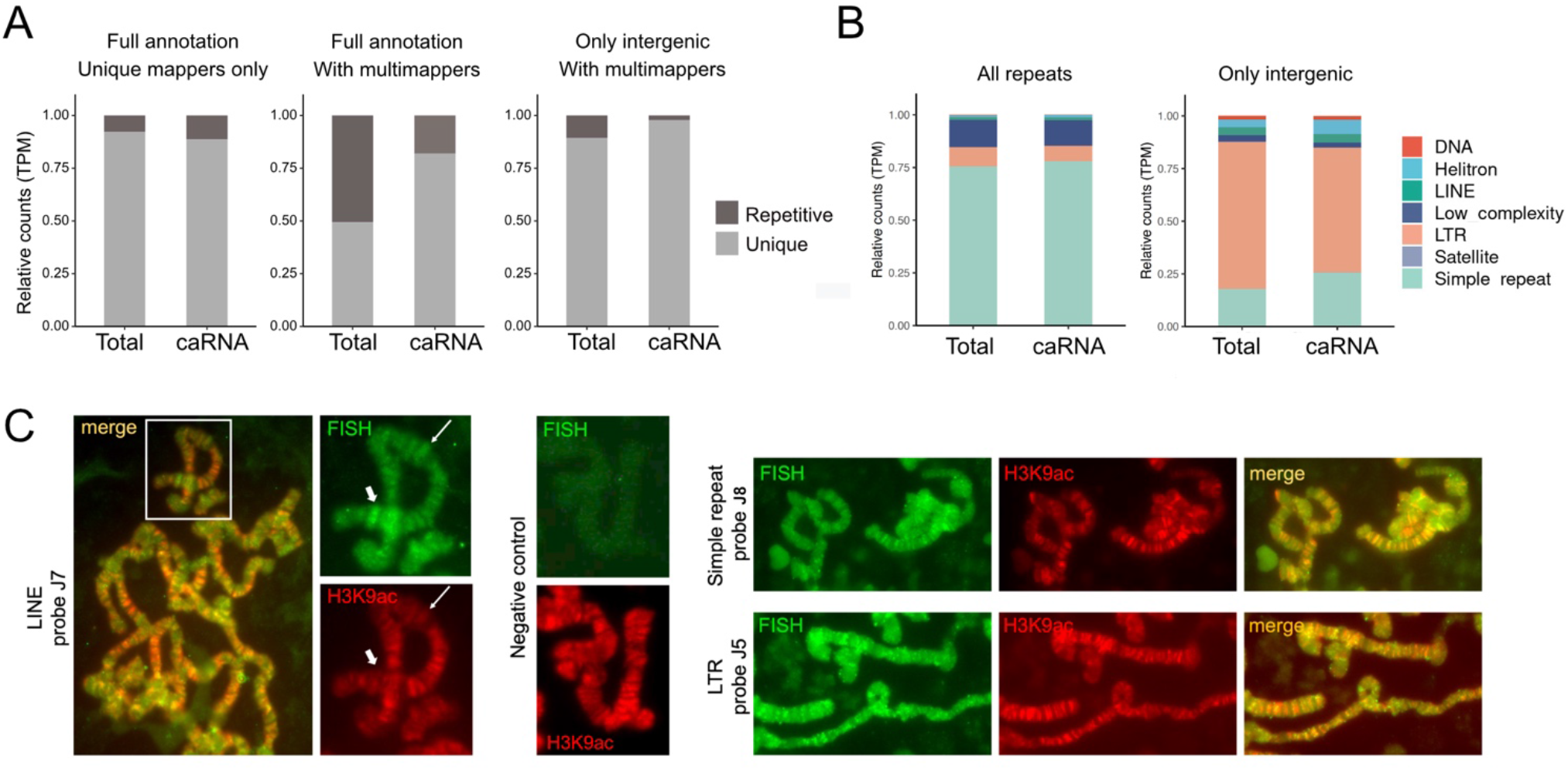
Identification of repetitive caRNAs (rep-caRNAs) in S2 cells. **(A)** The bar plots show the relative normalized transcript abundances (TPM) for unique and repetitive transcripts in total and chromatin RNA preparations. Each panel contains shows a different read counting strategy for rep-caRNA analysis. Left: uniquely mapping reads only. Middle: including reads mapping to multiple loci. Right: including reads mapping to multiple loci but excluding transcripts that overlap with annotated genes. Chromatin fraction normalized RNA abundance was averaged across replicates. **(B)** Relative distribution of normalized transcript abundances (TPM) per repetitive family. Left panel includes all the repetitive elements and right panel contains only intergenic repetitive elements. Total (left) and chromatin fraction (right) RNA preparations are depicted on each panel. Chromatin fraction normalized RNA abundance was averaged across replicates. **(C)** Fluorescence in-situ hybridization (FISH) showing the widespread distribution of three different rep-caRNAs in the salivary gland polytene chromosomes of *Drosophila melanogaster* (green). The preparations were counterstained with an antibody against H3K9ac (red). Magnification bars represent approximately 10 µm.

In order to ascribe each rep-caRNA read to a specific genomic coordinate, the repetitive annotation was extended by 50 bp both upstream and downstream of the annotated repeat, and only reads that mapped to a single location were included in the analysis. We could then detect rep-caRNAs expressed from 13 323 different repetitive loci in control cells (Table 2). Many of them were expressed at very low levels and were not detected in all replicate samples, which raises doubts about their biological significance, but as many as 3 249 were consistently detected with relatively high expression levels (at least 3 TPMs in each replicate, n=3).

A classification of all detected rep-caRNAs according to repeat families revealed that simple repeats and low complexity repeats were the most abundant rep-caRNA families. Intergenic rep-caRNAs were mainly derived from LTR elements (Fig. 4B). We carried out RNA-FISH in larval polytene chromosomes with probes complementary to three selected repeats to confirm that the rep-caRNAs detected by RNA-seq were associated to chromatin and widely distributed in the genome (Fig. 4C). The three probes hybridized to a large number of bands in the chromosomes arms and showed partial overlap with anti-H3K9ac staining, a mark for open chromatin.

In summary, many RNAs originating from specific repeat families -mainly simple repeats, low complexity repeats and LTRs - are associated with the chromatin. Although these rep-caRNAs constitute a minor fraction of the total chromatin-associated transcriptome, transcripts from more than 3000 different repeat loci could be consistently detected in the chromatin of control S2 cells.

### The exosome regulates the repetitive chromatin-associated transcriptome

Next we analyzed the effect of the exosome depletion on the abundance of rep-caRNAs using the 50 bp-extended annotation described above. As for unique sequences, the RRP6-kd and Double-kd gave stronger effects than the DIS3-kd, which was more similar to control cells as shown by both Spearman correlation and dimensionality reduction analysis (Suppl. Fig. S4B and S5B).

The number of expressed rep-caRNA loci that could be detected increased by approximately 50% in RRP6-kd and Double-kd (Table II). Some of these exosome-sensitive repeats were expressed at very low levels, but even if only high confidence transcripts were considered, the number of detected rep-caRNAs was almost twice as high in exosome depleted cells than in control cells (Table 2 and Suppl. Fig. S12). The rep-caRNAs that were revealed exclusively in exosome-depleted cells were mainly simple repeats (Suppl. Table S9).

Differential expression analysis identified 544 increased and 112 decreased rep-caRNAs in Double-kd, and all the repetitive families were increased (Fig. 5A-C, Table 1 and Suppl. Fig. S13). The number of differentially expressed rep-caRNAs was relatively low considering that rep-caRNAs from thousands of loci that were silent in control cells were revealed in Double-kd (compare Tables 1 and 2). This apparent discrepancy is explained by the fact that many rep-caRNAs detected upon exosome depletion are expressed at very low levels and differential expression algorithms give more confidence to robustly expressed transcripts. Interestingly, a direct comparison of the abundances of rep-caRNAs in control and knock-down samples including multimapping reads revealed that exosome depletion caused a global increase in the levels of low-abundance rep-caRNAs (Fig. 5D). Taken together, these results show that the exosome plays a key role in restricting the accumulation of repetitive RNAs in the chromatin, a role that is underestimated by conventional differential expression algorithms due to the low expression levels of many rep-caRNAs.

**Figure 5.**
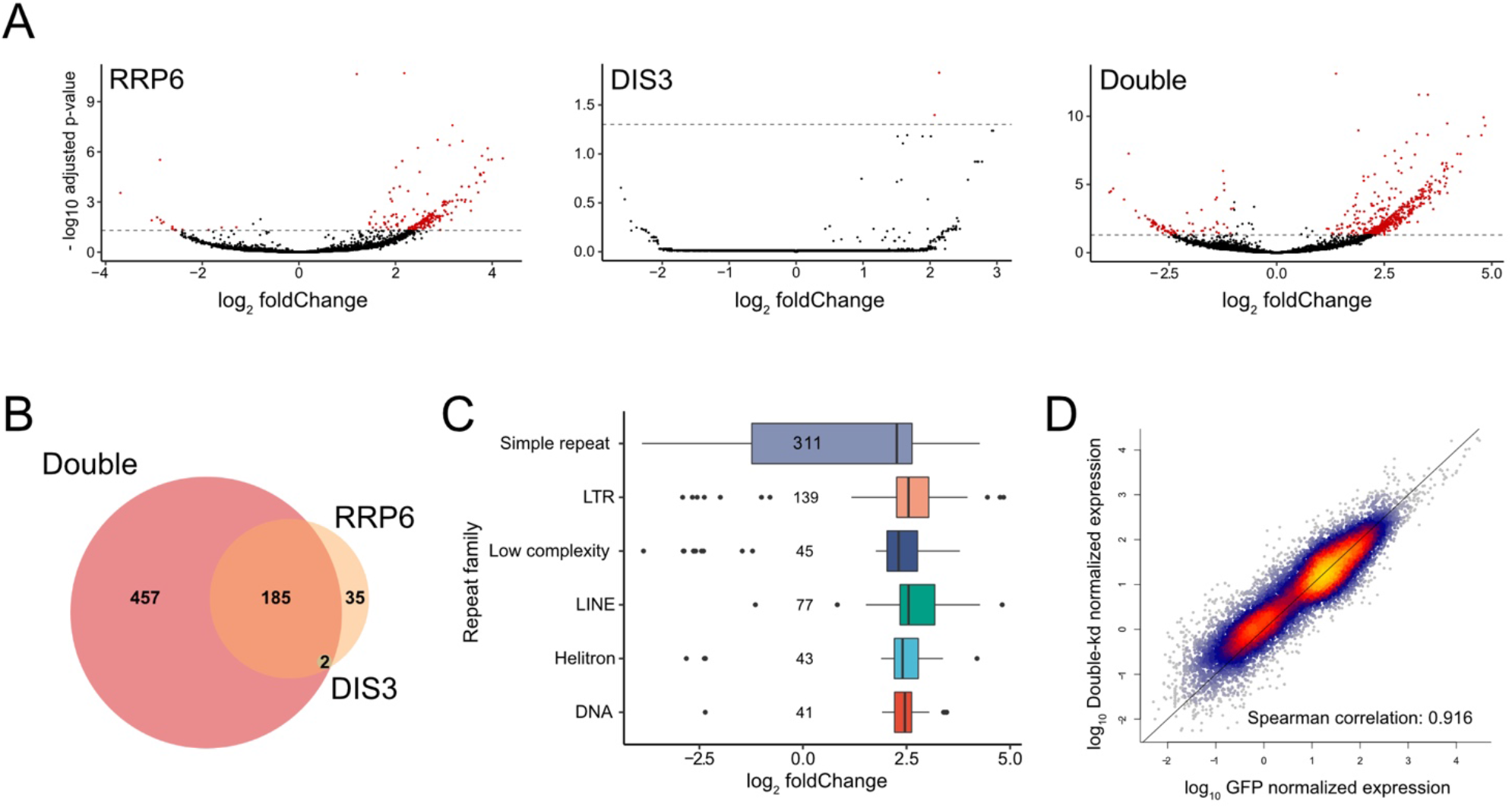
The effects of exosome depletion on the repetitive chromatin-associated transcriptome of S2 cells. **(A)** Volcano plots showing the results of rep-caRNAs differential expression analysis in Rrp6-kd, Dis3-kd or Double-kd versus control S2 cells. The x-axis represents the average RNA abundance change (log_2_ fold change). The y-axis represents the inverted log_10_(adjusted p-value). The dotted horizontal line corresponds to the cutoff significance value for differential expression (adjusted p-value < 0.05). Each dot represents a rep-caRNA transcript (n=16315). Significantly differentially expressed rep-caRNAs are denoted by red dots. The numbers of DE rep-caRNAs in each condition are provided in Table 1. **(B)** Intersection of significantly DE rep-caRNAs detected by differential expression analysis in the three depletion conditions. (p-value adjusted < 0.05). **(C)** Box plot showing the number of DE rep-caRNAs in each repeat family and the corresponding average RNA change (log_2_ fold change) in Double-kd versus control cells. The numbers correspond to the total number of DE rep-caRNAs in each repetitive family. The black lines inside the boxes denote the median for each specific repetitive family. **(D)** Scatter plot showing the normalized log-transformed rep-caRNA abundance (log_10_TPM) in Double-kd (y-axis) and control cells (x-axis). Rep-caRNA abundance was counted including multi-mapping reads. The dot density is color coded (yellow indicates higher transcript density, blue lower density). The RNA abundance correlation of double-kd and control conditions was tested using the Spearman correlation (correlation coeficient = 0.916). Black continuous line denotes the diagonal (perfect fit).

We asked whether the regulation of rep-caRNAs by the exosome was associated with specific chromatin environments and intersected the DE rep-caRNA loci with the nine chromatin states (state_1 to state_9) determined by [29] to investigate whether the regulation of rep-caRNA levels by the exosome was associated with specific chromatin environments. The majority of repeats included in the RepeatMasker annotation intersected with state_9, which corresponds to transcriptionally silent chromatin. The DE rep-caRNAs were instead strongly enriched in state_3 (intronic sequences, enhancer-like regions, regulatory regions) and state_7 (pericentromeric heterochromatin). Notably, polycomb-repressed regions (state_6) were underrepresented among DE rep-caRNAs (Fig. 6A). We also analyzed the chromosomal distribution of the DE rep-caRNAs (Fig. 6B), which revealed that the upregulated rep-caRNA loci could be classified into two major groups. One group, comprising 318 out of 544 upregulated rep-caRNAs, was mainly composed of simple repeats located in the chromosome arms. The second group, comprising 226 out of 544, included retrotransposons, mainly LTRs, located in the pericentromeric regions of chromosome arms 2L, 2R and 3L. A meta-analysis of publicly available ChIP-seq data showed that the DE rep-caRNAs of the simple repeat family located in the chromosome arms were significantly enriched in histone modifications such as H3K27ac and H3K4me1, which are marks characteristic of transcriptionally active regions, whereas the pericentromeric DE LTR-type rep-caRNAs had typical heterochromatin marks such as HP1 and H3K9me3 (Fig. 6C). In summary, the exosome regulates the repetitive chromatin-associated transcriptome and acts predominantly on two types of rep-caRNAs: simple repeat RNAs transcribed from active genic sequences and retrotransposon transcripts from the pericentromeric heterochromatin.

**Figure 6.**
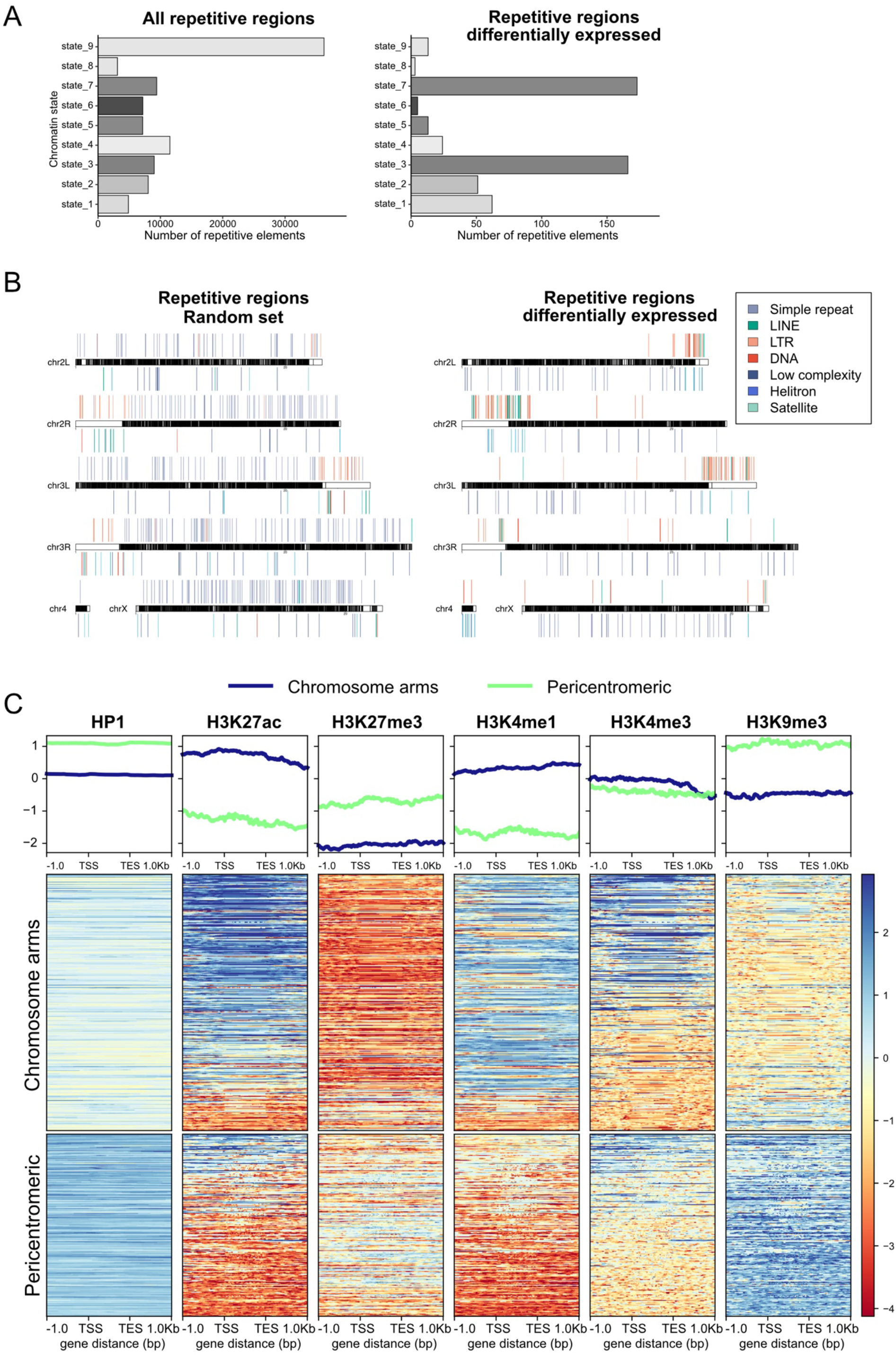
Characterization of the differentially expressed rep-caRNA loci in cells depleted of RRP6 and DIS3. **(A)** Intersection of DE rep-caRNAs with chromatin states [29]. The length of the bars represent the number of DE rep-caRNAs (x-axis) per chromatin state (y-axis). Left panel contains the entire rep-caRNA annotation. Right panel contains DE rep-caRNAs. State_1: active promoters and TSSs, state_2: transcriptional elongation and exons, state_3: intronic regions and regulatory regions, state_4: open chromatin, state_5: chromosome X and dosage compensation, state_6 CS6: polycomb repressed regions, state_7: pericentromeric heterochromatin, state_8: heterochromatin-like regions, state_9: silent chromatin. **(B)** Plot showing the genomic distribution of differentially expressed rep-caRNAs (left panel) and a random set of repetitive elements (right panel). The locations of rep-caRNA are indicated above and below the chromosome plots. Repeat families are color-coded equally as in A. The different *Drosophila melanogaster* <u>chromosomes</u> are depicted as black bars. White bands along the chromosomes represent citological bands. **(C)** Characterization of the chromatin environment of DE rep-caRNAs. The figure shows metagenes (top) and heatmaps (bottom) of publicly available ChIP-seq data of histone modifications and HP1 (accession numbers available in Suppl. Table 5). The metagenes show the average enrichment for each histone modification in pericentromeric DE rep-caRNAs (light green) and chromosome arms DE rep-caRNAs. The heatmap shows enrichment for each specific transcript. Enrichment in heatmap is color coded, where blue means enrichment and red depletion (color bar on the right). Coverage was normalized by library depth before computing enrichment.

### The exosome is required to maintain the packaging of the pericentromeric chromatin

Exosome depletion not only affected rep-caRNA levels but also the packaging of the repetitive chromatin. As many as 64% of the DARs (3281 out of 5105) overlapped with annotated repeats, many of them located within protein-coding genes (Fig. 7A). Moreover, in Double-kd, the chromatin of DE rep-caRNA loci was more accessible than that of a random set of rep-caRNA loci (Fig. 7B), which again suggests that RNA accumulation in the chromatin caused by exosome depletion leads to increased chromatin accessibility.

**Figure 7.**
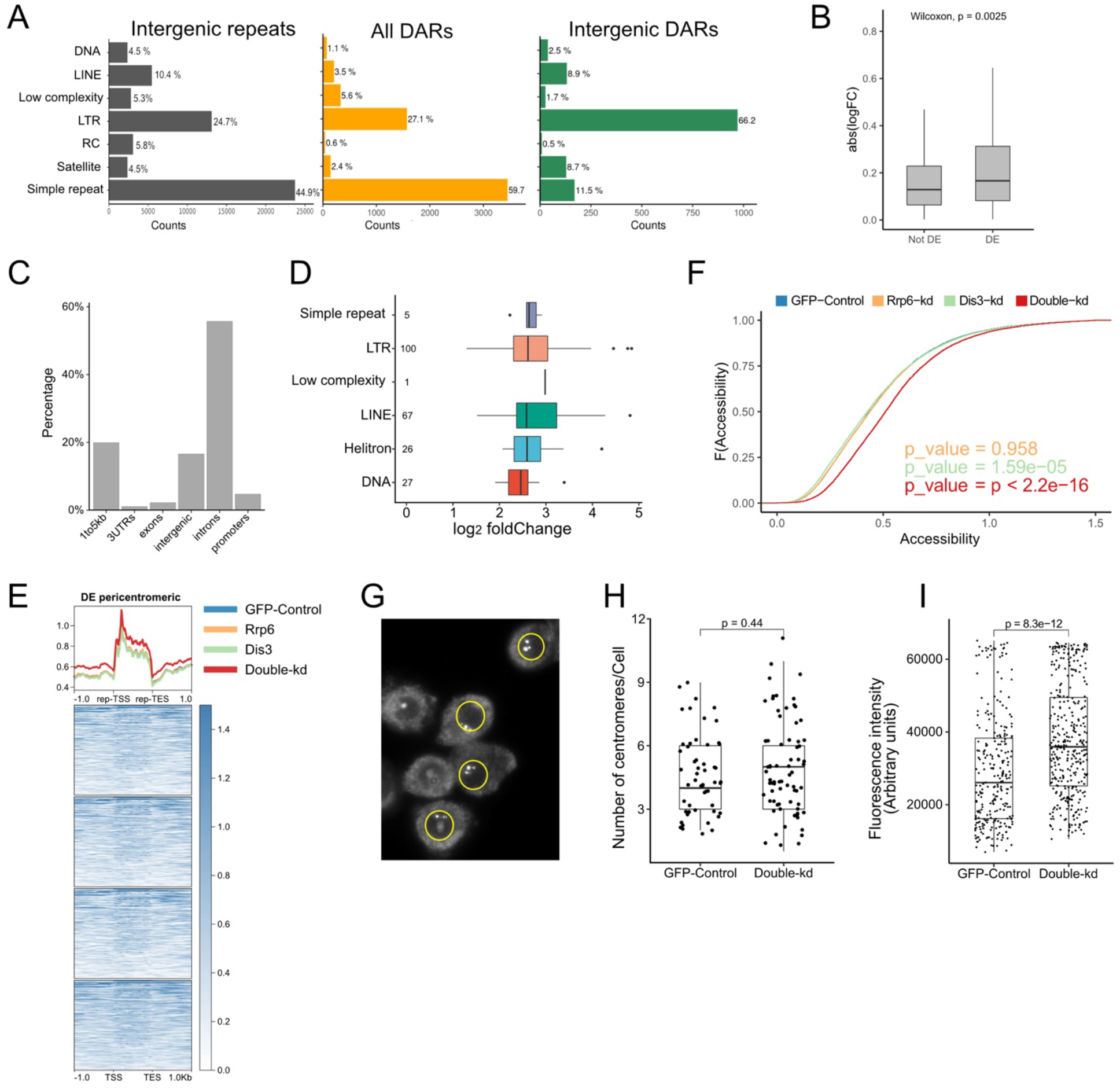
Depletion of RRP6 and DIS3 disrupts the packaging of the pericentromeric chromatin. **(A)** Association of DARs with specific repeat families. The left plot (dark grey) shows the fraction of intergenic repeats corresponding to each repeat family, for reference. The middle plot (yellow) shows the distribution of DARs in the different repeat families expressed as percentage of DARs belonging to each family. The right plot (green) shows the distribution of intergenic DARs in the different repeat families. **(B)** The box plot shows the absolute change in chromatin accessibility (y-axis, | log_2_FoldChange |) in DE rep-caRNA loci (left box) and in a random rep-caRNA control set expressed (x-axis). Black line inside the box denotes the median chromatin compaction change for each group. Difference between DE rep-caRNAs and control set was tested with a two-sided non-parametric Wilcoxon test (p-value shown in the plot). **(C)** Box plot showing the distribution of the average RNA change (log_2_ fold change) in double-kd versus control cells for increaseed pericentromeric rep-caRNAs. Each repetitive repetitive family is indicated in a different box. Numbers correspond to the total number of DE rep-caRNAs per repetitive family. Black line inside the box denotes the median for each specific repetitive family. **(D)** The bar plot shows the percentage of increased pericentromeric DE rep-caRNAs with annotated gene features (x-axis). **(E)** Chromatin compaction profiles of normalized average ATAC-seq signal across the DE pericentromeric rep-caRNAs. The figure shows the average compaction (metagene, top) and individual compaction values (heatmap, bottom) of chromatin accessibility in control cells (dark blue line) and in the different RNA exosome knockdown conditions (RRP6, blue; DIS3, green; and double knockdown, orange) in the increased pericentromeric rep-caRNAs. Compaction coverage includes the upstream and downstream 1 kb flanking regions. Signal has been subjected to sequencing depth normalization. **(F)** Chromatin compaction profiles of normalized average ATAC-seq signal across the entire pericentromeric region in control cells (dark blue) and in the different RNA exosome knockdown conditions (RRP6, blue; DIS3, green; and double knockdown, orange). The pericentromeric region [54] was divided into 1-kb bins and average compaction signal profiles were computed for the entire pericentromeric region. Signal has been subjected to sequencing depth normalization. **(G)** Immunofluorescence of CID protein in control S2 cells. The figure shows an example of a representative image used for quantification. The centromeres appear as bright dots inside the nucleus. Cell nuclei encircled in yellow are used as regions of interest for quantitative analysis of centromeric features. **(H)** Quantitative analysis of the number of centromeres. The box plots show the number of CID-positive foci per nucleus in control cells (left) and in cells depleted of both RRP6 and DIS3 (right). The black dots show the measures of individual cells. A two-sided t-test was used for comparison between conditions (n_GFP_ = 54, n_Double-kd_ = 73). **(I)** Quantitative analysis of centromere fluorescence intensity. The box plots show the fluorescence intensity of individual CID foci (y-axis, arbitrary units) in control cells (left) and in cells depleted of both RRP6 and DIS3 (right). Black dots correspond to individual foci measurements. Two-sided t-test was used for comparison between conditions (p-value < 0.0001).

A classification of the repetitive regions that intersected with DARs revealed a predominance of LTRs that was even more pronounced when only intergenic DARs were considered (Fig. 7A). Gypsy elements were particularly abundant in this group and constituted almost 85% of the LTRs that intersected with DARs, which revealed the role of the exosome in controlling this type of transposons.

The LTRs that were deregulated in Double-kd were mainly located in the pericentromeric regions of chromosomes 2 and 3, as shown in Fig. 6B. Pericentromeric rep-caRNA increased in exosome depleted cells were preferentially located in introns of protein-coding genes (Fig. 7C) and included LTRs, LINEs, Helitrons and other DNA transposons (Fig. 7D). A metagene analysis of ATAC-seq data showed that, in control cells, the chromatin of the DE pericentromeric rep-caRNA loci was more accessible than that of their surrounding sequences. Moreover, the chromatin accessibility of the DE pericentromeric rep-caRNA loci was significantly higher in Double-kd than in control (Fig. 7E), in agreement with the observation that many DARs overlapped with DE rep-caRNA loci. However, the increased chromatin accessibility was not restricted to the rep-caRNA loci but affected also the rep-caRNA flanking regions, which suggested that exosome depletion had a global effect on pericentromeric chromatin compaction. A cumulative plot of chromatin accessibility frequencies across the entire pericentromeric region confirmed that this was the case (Fig. 7F). Based on this observation, we asked whether the exosome depletion had any morphologically detectable effect on the overall organization of the centromeres. We visualized the centromeres of S2 cells by immunofluorescence using an antibody against CID, a centromere-specific histone H3 variant that functions as an epigenetic mark for centromere identity (Fig. 7G). The number of centromeres per cell was not affected by the exosome depletion (Fig. 7H), but the average fluorescence intensity of the centromeres was significantly higher in exosome depleted cells (Fig. 7I). We could not establish whether the increased fluorescence intensity was due to increased CID incorporation into the centromeres or to more efficient CID detection due to increased chromatin accessibility. In any case, the increased fluorescence intensity showed that exosome depletion results in a widespread decompaction of pericentromeric chromatin with a concomitant alteration of centromere architecture.

## DISCUSSION

The RNA exosome is bound to chromosomes and to transcribed genes, as shown by immunofluorescence and ChIP experiments [25,36,37], and it has been assumed that RNA degradation by the exosome takes place, at least to some extent, in the chromatin [41]. Here we have characterized the chromatin-associated transcriptome to directly address the role of the RNA exosome in the chromatin. We have shown that the chromatin has a specific RNA composition, we have identified different types of transcripts that are targeted by the exosome in the chromatin, and our results reveal that the exosome not only restricts caRNA levels, but also influences chromatin accessibility and gene regulation. A relevant question in the context of the present study is the nature of the caRNAs. Some of them represent nascent transcripts whereas others are likely RNAs that remain bound to the chromatin after transcription, perhaps because they play a role in the chromatin or because they lack efficient export mechanisms and are somehow protected from degradation. The abundances of the unique caRNAs that we detected under control conditions were relatively well correlated with the abundances of the corresponding sequences in total RNA, which supports the view that unique caRNAs represent to a large extent nascent transcripts and that their abundance reflects the level of expression of the gene. The chromatin levels of most rep-caRNAs were instead poorly correlated with their total RNA levels, which suggests that rep-caRNAs constitute a more heterogeneous group of transcripts.

We knocked down each of the two exosome catalytic subunits separately to get insight into their specific contributions to caRNA degradation. The depletion of DIS3 had a minor effect, and our results suggest that RRP6 is the main responsible for the degradation of caRNAs in *Drosophila*. However, the double depletion of both RRP6 and DIS3 gave a more severe effect than the depletion of only RRP6, which implies that DIS3 is not totally dispensable in the chromatin and suggests the existence of two major types of caRNA exosome substrates. One type represents caRNAs that can be degraded by either DIS3 or RRP6 and are therefore only differentially expressed when both subunits are knocked down simultaneously. A second type includes caRNAs that require RRP6 for degradation and thus appear as differentially expressed in both the RRP6-kd and Double-kd conditions.

Depletion of exosome subunits by RNAi has both direct and indirect effects on the transcriptome. We have addressed this question by using ChIP-seq data for RRP6, which has allowed us to establish the presence of the exosome at the affected loci. In this way, we could discriminate between different types of exosome-sensitive genes. RRP6 occupancy was low in approximately 60% of the exosome-sensitive genes and we favor the conclusion that the changes observed at these loci were secondary effects of altered transcription regulation. The remaining 40% of the exosome-sensitive genes showed high RRP6 occupancy and are likely to be direct exosome targets. Some of the direct targets encode regulatory factors with roles in cell differentiation and morphogenesis, and changes in their expression can explain the deregulation observed at loci with low RRP6 occupancy.

The exosome-sensitive genes that showed altered chromatin accessibility in exosome-depleted cells, referred to as DEG_DARs in our study, were characterized by high RRP6 occupancy and were bound by both Polycomb and trithorax factors. Polycomb repressed genes are typically silenced and show high levels of H3K27me3, but Polycomb factors have also been detected in active loci that show a “balanced” chromatin state characterized by low H3K27me3 and by the simultaneous presence of Polycomb and trithorax factors (see [40] and references therein). It has been proposed that these balanced chromatin states can become fully activated or fully repressed during development, but the dynamics of their regulation is not fully understood. Our results suggest that local RNA clearance by the exosome is a key component in this regulatory circuit and we propose that caRNA degradation by the exosome at these loci is required to avoid the accumulation of RNAs that would destabilize the local chromatin homeostasis by sequestering Polycomb factors through a process that would resemble that proposed by Garland et al [27] in mammalian cells. The physical interactions detected between exosome subunits and Su(z)12, dRING and Trl/GAF provide a possible mechanism for the recruitment of the exosome to the “balanced” loci.

Approximately 35% of the *Drosophila melanogaster* genome is composed of repetitive sequences the expression of which has been poorly characterized [42,43]. We addressed the study of the repetitive chromatin-associated transcriptome because of its potential impact on chromosome architecture and homeostasis. Although the analysis of repetitive sequences is constrained by technical limitations, we have been able to draw conclusions about the overall abundance of rep-caRNA families and about the role of the exosome in controlling rep-caRNA levels. We have detected rep-caRNAs produced from approximately 2.5 % of the annotated repeats. Repetitive RNAs stably associated with the chromatin have also been reported in mammalian cells [44], which suggests that the presence of repetitive RNA in the chromatin is a general feature of metazoans.

The fact that the exosome degrades specific repetitive RNAs has been previously reported. These include noncoding RNAs derived from tandemly repeated elements at an *Hsp70* locus in *D. melanogaster* [45], human endogenous retroviral transcripts [46] and transcribed repeat RNAs that are produced as a result of nucleotide repeat expansions in the C9orf72 gene in human cells [47]. Moreover, small-RNA profiling of rrp6 null *D. melanogaster* larvae revealed an increase in reads mapping to repeat elements [48] and we have previously shown that RRP6 silences a subset of transposons and heterochromatic repeats in S2 cells [25]. Our present study extends these earlier observations and reveals that the degradation of many repetitive RNAs by the exosome takes place in the chromatin, which is particularly relevant due to the fact that RNA can compromise the packaging of the chromatin through different mechanisms. RNA can interact with chromatin proteins such as HP1 and induce their dissociation from the H3K9-methylated histone tails [22,25], reduce the electrostatic interactions between the histones tails and the DNA [19], engage in the formation of R-loops that promote chromatin decondensation [49] or direct the assembly of biomolecular condensates through liquid-liquid phase-separation phenomena [50,51]. Controlling caRNA levels is thus a necessary mechanisms of chromatin homeostasis and our ATAC-seq experiments show that the exosome is required for the maintenance of chromatin packaging. This role is particularly relevant in the pericentromeric regions where exosome depletion results into increased levels of pericentromeric LTR transcripts, global chromatin unfolding and altered centromere organization.

The repetitive RNAs that we have identified as exosome targets can be ascribed to different types of chromatin environments and differ in a series of features such as chromosomal location, level of expression and histone marks, but they all share a common trait: low levels of H3K27me3. This observation suggests that the exosome acts in chromatin regions that are not silenced by the Polycomb complexes. A possible explanation is that Polycomb repression is effective enough to prevent pervasive transcription. However, there is experimental evidence involving ncRNAs in Polycomb regulation (reviewed in [52]) and studies in *C. elegans* have revealed that another exoribonuclease, XRN2, targets transcripts from H3K27me3-silenced loci and contributes to maintain Polycomb loci repressed [26]. Yet a third ribonuclease complex, Ccr4-Not, has been shown to degrade chromatin-associated telomeric transcripts in the *Drosophila* germline [53]. Altogether, these findings suggest that RNA degradation in the chromatin is a widespread phenomenon and that each chromatin environment recruits specific ribonucleases, probably through specialized protein-protein interactions, to control the levels of different types of chromatin-associated RNAs.

## ACKNOWLEDGEMENTS

We thank Alisa Alekseenko (SciLifeLab, Karolinska Institute) for assistance with Illumina sequencing and Jakub Westholm (NBIS, SciLifeLab, Stockholm University) for bioinformatics support. The computations were enabled by resources in project [SNIC 2018/8-294] provided by the Swedish National Infrastructure for Computing (SNIC) at UPPMAX, partially funded by the Swedish Research Council through grant agreement no. 2018-05973. We thank the Imaging Facility at Stockholm University (IFSU) for support with microscopy.

## REFERENCES

1. Li,X. and Fu,X.D. (2019) Chromatin-associated RNAs as facilitators of functional genomic interactions. Nat. Rev. Genet. 2019 209, 20, 503–519.

2. Meller,V.H., Joshi,S.S. and Deshpande,N. (2015) Modulation of Chromatin by Noncoding RNA. https://doi.org/10.1146/annurev-genet-112414-055205, 49, 673–695.

3. Brockdorff,N., Bowness,J.S. and Wei,G. (2020) Progress toward understanding chromosome silencing by Xist RNA. Genes Dev., 34, 733–744.

4. Volpe,T.A., Kidner,C., Hall,I.M., Teng,G., Grewal,S.I.S. and Martienssen,R.A. (2002) Regulation of heterochromatic silencing and histone H3 lysine-9 methylation by RNAi. Science (80-.)., 297, 1833–1837.

5. Ozata,D.M., Gainetdinov,I., Zoch,A., O’Carroll,D. and Zamore,P.D. (2019) PIWI-interacting RNAs: small RNAs with big functions. Nat. Rev. Genet., 20, 89–108.

6. Statello,L., Guo,C.J., Chen,L.L. and Huarte,M. (2021) Gene regulation by long non-coding RNAs and its biological functions. Nat. Rev. Mol. Cell Biol., 22, 96–118.

7. Tippens,N.D., Vihervaara,A. and Lis,J.T. (2018) Enhancer transcription: what, where, when, and why? Genes Dev., 32, 1–3.

8. Belostotsky,D. (2009) Exosome complex and pervasive transcription in eukaryotic genomes. Curr. Opin. Cell Biol., 21, 352–358.

9. Rougemaille,M. and Libri,D. (2010) Control of cryptic transcription in eukaryotes. Adv. Exp. Med. Biol., 702, 122–131.

10. Bonath,F., Domingo-Prim,J., Tarbier,M., Friedländer,M.R. and Visa,N. (2018) Next-generation sequencing reveals two populations of damage-induced small RNAs at endogenous DNA doublestrand breaks. Nucleic Acids Res., 46, 11869–11882.

11. Januszyk,K. and Lima,C.D. (2014) The eukaryotic RNA exosome. Curr. Opin. Struct. Biol., 24, 132–140.

12. Lebreton,A., Tomecki,R., Dziembowski,A. and Séraphin,B. (2008) Endonucleolytic RNA cleavage by a eukaryotic exosome. Nat, 456, 993–996.

13. Chlebowski, A., Lubas, M., Jensen, T. H. and Dziembowski, A. (2013) RNA decay machines: The exosome. Biochim. Biophys. Acta - Gene Regul. Mech. 1829, 552–560.

14. Kilchert,C., Wittmann,S. and Vasiljeva,L. (2016) The regulation and functions of the nuclear RNA exosome complex. Nat. Rev. Mol. Cell Biol. 2016 174, 17, 227–239.

15. Domingo-Prim,J., Endara-Coll,M., Bonath,F., Jimeno,S., Prados-Carvajal,R., Friedländer,M.R., Huertas,P. and Visa,N. (2019) EXOSC10 is required for RPA assembly and controlled DNA end resection at DNA double-strand breaks. Nat. Commun., 10.

16. Davidson,L., Francis,L., Cordiner,R.A., Eaton,J.D., Estell,C., Macias,S., Cáceres,J.F. and West,S. (2019) Rapid Depletion of DIS3, EXOSC10, or XRN2 Reveals the Immediate Impact of Exoribonucleolysis on Nuclear RNA Metabolism and Transcriptional Control. Cell Rep., 26, 2779-2791.e5.

17. Kiss,D.L. and Andrulis,E.D. (2010) Genome-wide analysis reveals distinct substrate specificities of Rrp6, Dis3, and core exosome subunits. RNA, 16, 781–791.

18. Wu,M., Karadoulama,E., Lloret-Llinares,M., Rouviere,J.O., Vaagensø,C.S., Moravec,M., Li,B., Wang,J., Wu,G., Gockert,M., et al. (2020) The RNA exosome shapes the expression of key protein-coding genes. Nucleic Acids Res., 48, 8509.

19. Dueva,R., Akopyan,K., Pederiva,C., Trevisan,D., Dhanjal,S., Lindqvist,A. and Farnebo,M. (2019) Neutralization of the Positive Charges on Histone Tails by RNA Promotes an Open Chromatin Structure. Cell Chem. Biol., 26, 1436–1449.e5.

20. Hendrickson,D., Kelley,D.R., Tenen,D., Bernstein,B. and Rinn,J.L. (2016) Widespread RNA binding by chromatin-associated proteins. Genome Biol., 17, 1–18.

21. He,C., Sidoli,S., Warneford-Thomson,R., Tatomer,D.C., Wilusz,J.E., Garcia,B.A. and Bonasio,R. (2016) High-Resolution Mapping of RNA-Binding Regions in the Nuclear Proteome of Embryonic Stem Cells. Mol. Cell, 64, 416–430.

22. Keller,C., Adaixo,R., Stunnenberg,R., Woolcock,K.J., Hiller,S. and Bühler,M. (2012) HP1 Swi6 Mediates the Recognition and Destruction of Heterochromatic RNA Transcripts. Mol. Cell, 47, 215–227.

23. Davidovich,C., Zheng,L., Goodrich,K.J. and Cech,T.R. (2013) Promiscuous RNA binding by Polycomb repressive complex 2. Nat. Struct. Mol. Biol., 20, 1250–1257.

24. Wang,X., Goodrich,K.J., Gooding,A.R., Naeem,H., Archer,S., Paucek,R.D., Youmans,D.T., Cech,T.R. and Davidovich,C. (2017) Targeting of Polycomb Repressive Complex 2 to RNA by Short Repeats of Consecutive Guanines. Mol. Cell, 65, 1056–1067.e5.

25. Eberle,A.B., Jordán-Pla,A., Gañez-Zapater,A., Hessle,V., Silberberg,G., von Euler,A., Silverstein,R.A. and Visa,N. (2015) An Interaction between RRP6 and SU(VAR)3-9 Targets RRP6 to Heterochromatin and Contributes to Heterochromatin Maintenance in Drosophila melanogaster. PLoS Genet., 11.

26. Mattout,A., Gaidatzis,D., Padeken,J., Schmid,C.D., Aeschimann,F., Kalck,V. and Gasser,S.M. (2020) LSM2-8 and XRN-2 contribute to the silencing of H3K27me3-marked genes through targeted RNA decay. Nat. Cell Biol., 22, 579–590.

27. Garland,W., Comet,I., Wu,M., Radzisheuskaya,A., Rib,L., Vitting-Seerup,K., Lloret-Llinares,M., Sandelin,A., Helin,K. and Jensen,T.H. (2019) A Functional Link between Nuclear RNA Decay and Transcriptional Control Mediated by the Polycomb Repressive Complex 2. Cell Rep., 29, 1800.

28. Eberle,A.B., Hessle,V., Helbig,R., Dantoft,W., Gimber,N. and Visa,N. (2010) Splice-Site Mutations Cause Rrp6-Mediated Nuclear Retention of the Unspliced RNAs and Transcriptional Down-Regulation of the Splicing-Defective Genes. PLoS One, 5, e11540.

29. Kharchenko,P. V., Alekseyenko,A.A., Schwartz,Y.B., Minoda,A., Riddle,N.C., Ernst,J., Sabo,P.J., Larschan,E., Gorchakov,A.A., Gu,T., et al. (2011) Comprehensive analysis of the chromatin landscape in Drosophila. Nature, 471, 480.

30. Buenrostro,J.D., Wu,B., Chang,H.Y. and Greenleaf,W.J. (2015) ATAC-seq: A Method for Assaying Chromatin Accessibility Genome-Wide. Curr. Protoc. Mol. Biol., 109, 21.29.1.

31. Yu,S., Jordán-Pla,A., Gañez-Zapater,A., Jain,S., Rolicka,A., Farrants,A.K.Ö. and Visa,N. (2018) SWI/SNF interacts with cleavage and polyadenylation factors and facilitates pre-mRNA 3′ end processing. Nucleic Acids Res., 46, 8557–8573.

32. Ewels,P.A., Peltzer,A., Fillinger,S., Patel,H., Alneberg,J., Wilm,A., Garcia,M.U., Di Tommaso,P. and Nahnsen,S. (2020) The nf-core framework for community-curated bioinformatics pipelines. Nat. Biotechnol. 2020 383, 38, 276–278.

33. Bentsen,M., Goymann,P., Schultheis,H., Klee,K., Petrova,A., Wiegandt,R., Fust,A., Preussner,J., Kuenne,C., Braun,T., et al. (2020) ATAC-seq footprinting unravels kinetics of transcription factor binding during zygotic genome activation. Nat. Commun. 2020 111, 11, 1–11.

34. Gramates,L.S., Marygold,S.J., Dos Santos,G., Urbano,J.M., Antonazzo,G., Matthews,B.B., Rey,A.J., Tabone,C.J., Crosby,M.A., Emmert,D.B., et al. (2017) FlyBase at 25: looking to the future. Nucleic Acids Res., 45, D663–D671.

35. Hessle,V., Björk,P., Sokolowski,M., De Valdivia,E.G., Silverstein,R., Artemenko,K., Tyagi,A., Maddalo,G., Ilag,L., Helbig,R., et al. (2009) The Exosome Associates Cotranscriptionally with the Nascent Pre-mRNP through Interactions with Heterogeneous Nuclear Ribonucleoproteins. Mol. Biol. Cell, 20, 3459.

36. Andrulis,E.D., Werner,J., Nazarian,A., Erdjument-Bromage,H., Tempst,P. and Lis,J.T. (2002) The RNA processing exosome is linked to elongating RNA polymerase II in Drosophila. Nature, 420, 837–841.

37. Lim,S.J., Boyle,P.J., Chinen,M., Dale,R.K. and Lei,E.P. (2013) Genome-wide localization of exosome components to active promoters and chromatin insulators in Drosophila. Nucleic Acids Res., 41, 2963–2980.

38. Schaaf,C.A., Misulovin,Z., Gause,M., Koenig,A., Gohara,D.W., Watson,A. and Dorsett,D. (2013) Cohesin and polycomb proteins functionally interact to control transcription at silenced and active genes. PLoS Genet., 9.

39. Frey,F., Sheahan,T., Finkl,K., Stoehr,G., Mann,M., Benda,C. and Müller,J. (2016) Molecular basis of PRC1 targeting to polycomb response elements by PhoRC. Genes Dev., 30, 1116–1127.

40. Schwartz,Y.B., Kahn,T.G., Stenberg,P., Ohno,K., Bourgon,R. and Pirrotta,V. (2010) Alternative Epigenetic Chromatin States of Polycomb Target Genes. PLOS Genet., 6, e1000805.

41. Eberle,A.B. and Visa,N. (2014) Quality control of mRNP biogenesis: networking at the transcription site. Semin. Cell Dev. Biol., 32, 37–46.

42. Hoskins,R.A., Carlson,J.W., Kennedy,C., Acevedo,D., Evans-Holm,M., Frise,E., Wan,K.H., Park,S., Mendez-Lago,M., Rossi,F., et al. (2007) Sequence Finishing and Mapping of Drosophila melanogaster Heterochromatin. Science, 316, 1625.

43. Krassovsky,K. and Henikoff,S. (2014) Distinct chromatin features characterize different classes of repeat sequences in Drosophila melanogaster. BMC Genomics, 15, 105.

44. Hall,L.L., Carone,D.M., Gomez,A. V., Kolpa,H.J., Byron,M., Mehta,N., Fackelmayer,F.O. and Lawrence,J.B. (2014) Stable C0T-1 repeat RNA is abundant and is associated with euchromatic interphase chromosomes. Cell, 156, 907–919.

45. Kuan,Y.-S., Brewer-Jensen,P., Bai,W.-L., Hunter,C., Wilson,C.B., Bass,S., Abernethy,J., Wing,J.S. and Searles,L.L. (2009) Drosophila Suppressor of Sable Protein [Su(s)] Promotes Degradation of Aberrant and Transposon-Derived RNAs. Mol. Cell. Biol., 29, 5590–5603.

46. Kammler,S., Lykke-Andersen,S. and Jensen,T.H. (2008) The RNA Exosome Component hRrp6 Is a Target for 5-Fluorouracil in Human Cells. Mol. Cancer Res., 6, 990–995.

47. Kawabe,Y., Mori,K., Yamashita,T., Gotoh,S. and Ikeda,M. (2020) The RNA exosome complex degrades expanded hexanucleotide repeat RNA in C9orf72 FTLD/ALS. EMBO J., 39.

48. Yamanaka,S., Mehta,S., Reyes-Turcu,F.E., Zhuang,F., Fuchs,R.T., Rong,Y., Robb,G.B. and Grewal,S.I.S. (2013) RNAi triggered by specialized machinery silences developmental genes and retrotransposons. Nature, 493, 557–560.

49. Chédin,F. (2016) Nascent connections: R-loops and chromatin patterning. Trends Genet., 32, 828–838.

50. Nozawa,R.S. and Gilbert,N. (2019) RNA: Nuclear Glue for Folding the Genome. Trends Cell Biol., 29, 201–211.

51. Rippe,K. (2022) Liquid-Liquid Phase Separation in Chromatin. Cold Spring Harb. Perspect. Biol., 14.

52. Almeida,M., Bowness,J.S. and Brockdorff,N. (2020) The many faces of Polycomb regulation by RNA. Curr. Opin. Genet. Dev., 61, 53–61.

53. Kordyukova,M., Sokolova,O., Morgunova,V., Ryazansky,S., Akulenko,N., Glukhov,S. and Kalmykova,A. (2020) Nuclear Ccr4-Not mediates the degradation of telomeric and transposon transcripts at chromatin in the Drosophila germline. Nucleic Acids Res., 48, 141–156.

54. Zenk,F., Zhan,Y., Kos,P., Löser,E., Atinbayeva,N., Schächtle,M., Tiana,G., Giorgetti,L. and Iovino,N. (2021) HP1 drives de novo 3D genome reorganization in early Drosophila embryos. Nat. 2021 5937858, 593, 289–293.

