## Supplementatry Figures for "The exosome degrades chromatin-associated RNAs genome-wide and maintains chromatin homeostasis"

### SUPPLEMENTARY FIGURES

Planells et al.

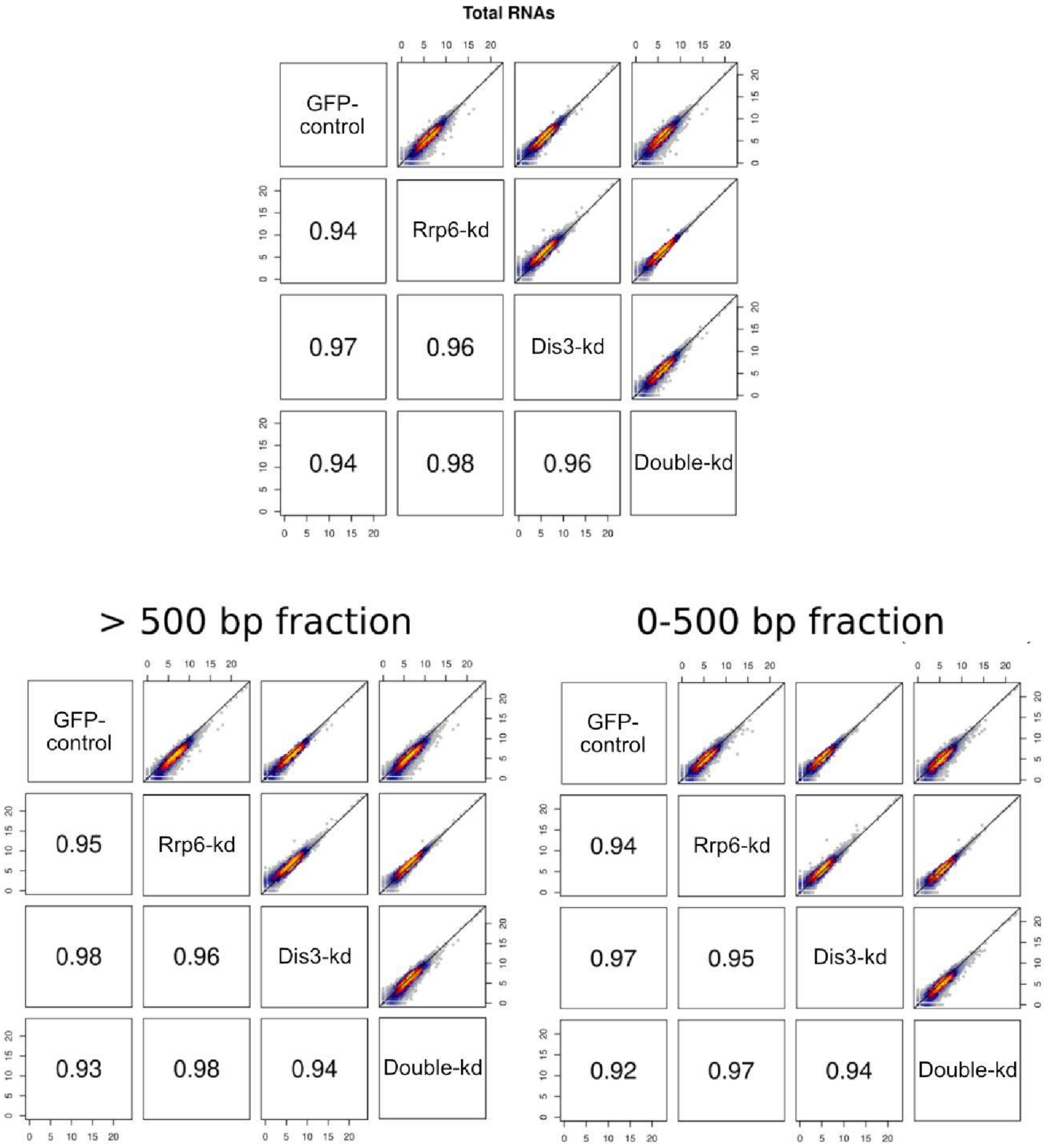


**Supplementary Figure S1. RNA-seq: correlations between biological replicates**. Pairwise comparisons of reads mapping to genes were used to compute the Spearman correlation coefficient. Each comparison is shown in the diagonal. The spearman correlation coefficients for the different comparisons are shown in the bottom-left half of each panel. Bin density is depicted in the top-right half of the panels. Yellow, red denote higher bin density while blue, gray show lower bin density. Top panel shows correlations for total RNA samples. Lower panels show correlations for the two different chromatin fractions described in material and methods. Biological replicates (n = 3) were merged at bam level with samtools merge function.

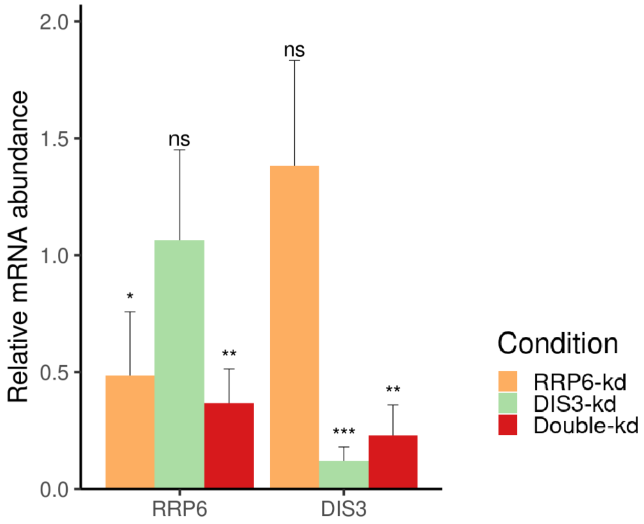


**Supplementary figure S2.** Efficiency of exosome depletion by RNAi. RRP6 and DIS3 mRNA levels measured by RT-qPCR, and normalized to Actin5C. Primers sequences are provided in Supplementary Table S. Bar height shows the average relative mRNA levels and error bars denote standard deviation of the mRNA abundance in the replicates (n = 3). Statistical significance was tested using a 2-tailed Student's t-test. * p>0.05, ** p>0.005, *** p>0.0005. ns, not significant.

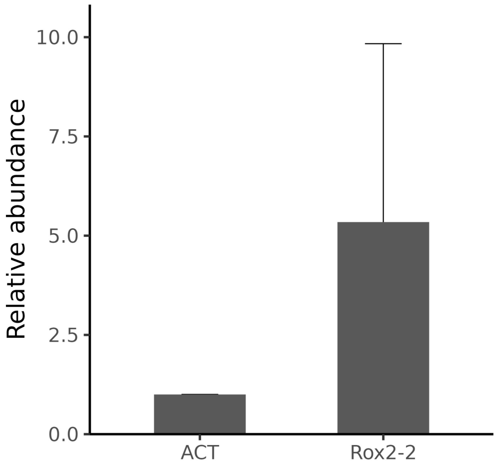


**Supplementary Figure S3. Enrichment of Rox2.2 lncRNA in the chromatin fraction**. The bars show the relative mRNA levels for Actin5C (ACT) and RoX2 transcripts quantified by RT-qPCR. The height of the bar shows the relative abundance of Actin5C (ACT) and RoX2 transcripts in chromatin RNA preparation versus total RNA preparation. Bars show averages and error bars correspond to standard deviations of mRNA and lncRNA abundance (n=4).

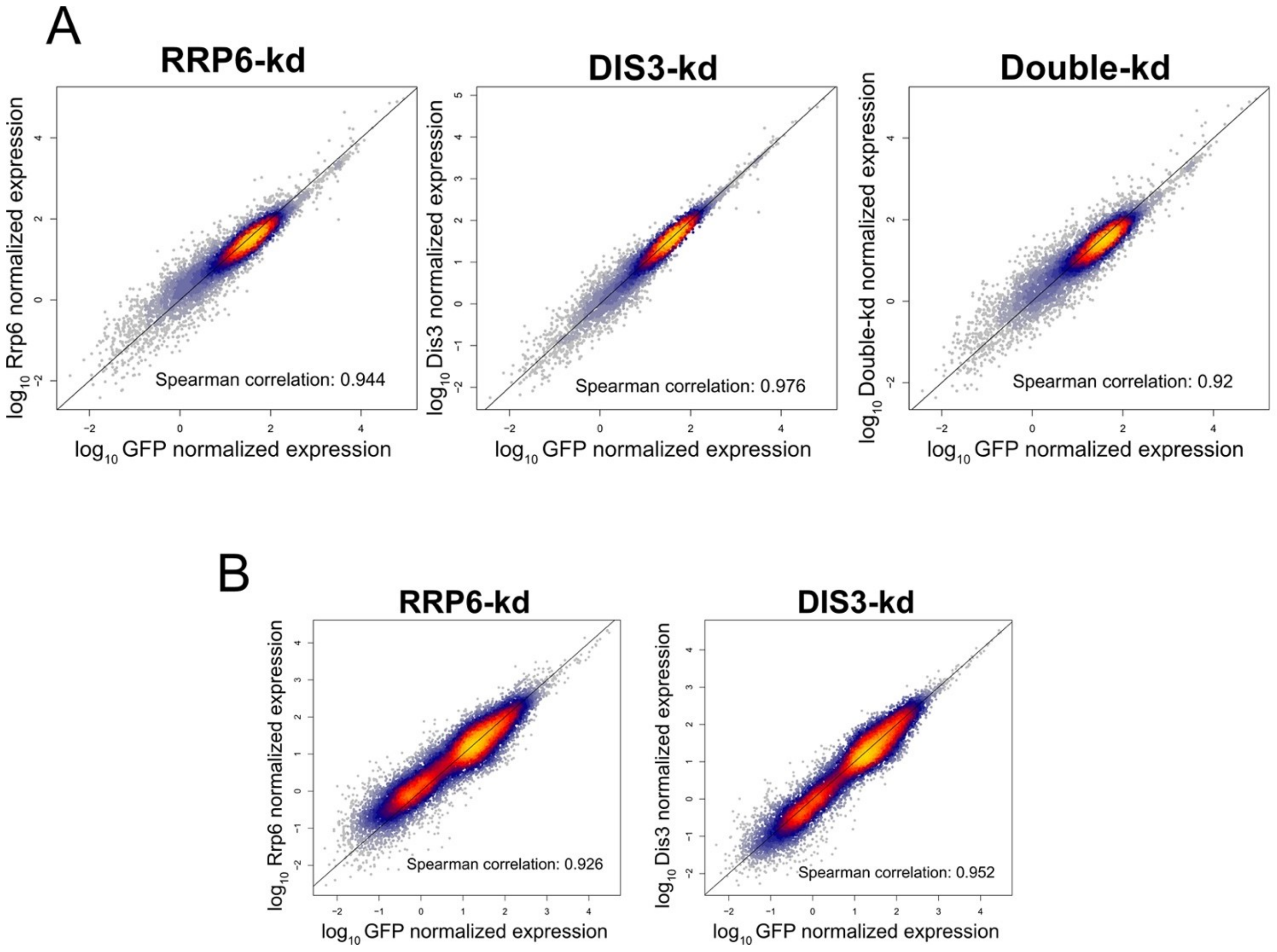


**Supplementary Figure S4. Correlation analysis of RNA-seq data in exosome depleted cells.** The scatter plots compare the average normalized log-transformed rep-caRNA abundance ($\log_{10}$TPM) in control (GFP) cells and RNA exosome depleted cells, as indicated. (**A**) shows data for unique reads only. (**B**) shows caRNA abundances including multimapping reads. Data for Double-kd in B is shown in Fig. 6D. The dot density is color coded (yellow indicates higher transcript density, grey lower density). Black continuous line denotes the diagonal (perfect fit).

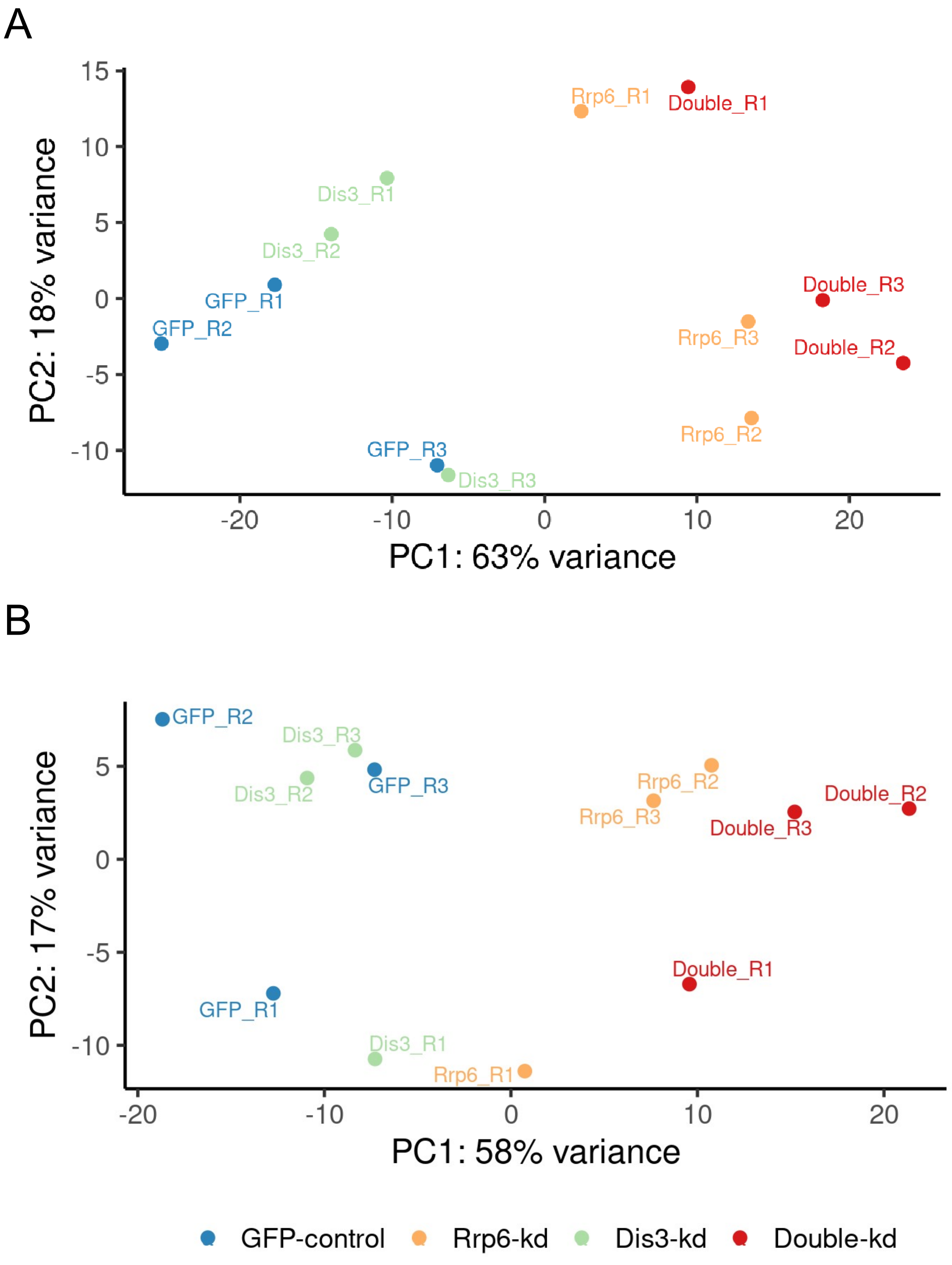


**Supplementary Figure S5. PCA analysis of RNA-seq data in exosome depleted cells.** Principal component analysis of unique **(A)** and repetitive **(B)** caRNA RNA-seq data. Principal component 1 (x-axes) separates the different conditions into two groups. DIS3 is similar to GFP control condition while RRP6 and the Double-kd are differentiated from the control condition. Transcript read counts have been transformed with the varianceStabilizingTranformation function contained in DESeq2 package.

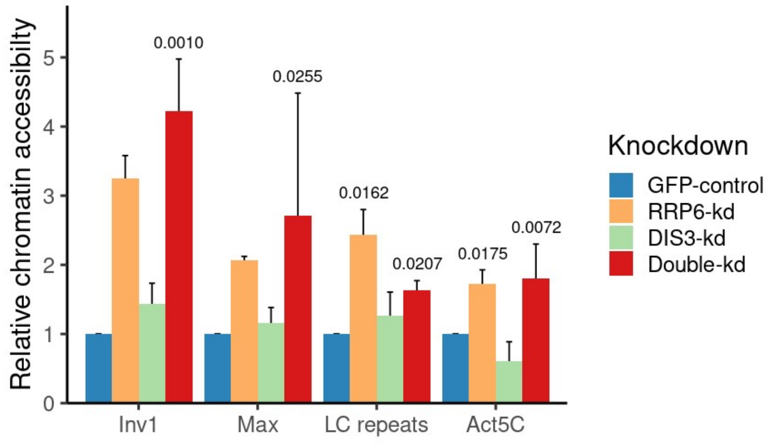


**Supplementary Figure S6. ATAC-qPCR analysis of rep-caRNA loci.** Chromatin accessibility was measured at four different caRNA loci by ATAC-qPCR. The bars shows the average chromatin accessibility (n = 3) and the error bars correspond to the standard deviation across replicates. Two-tailed student t-test was performed to test statistical significance among the observed differences compared to GFP-control. The PCR signal were normalized to Ctp. Primer sequences are provided in Suppl. Table S2.

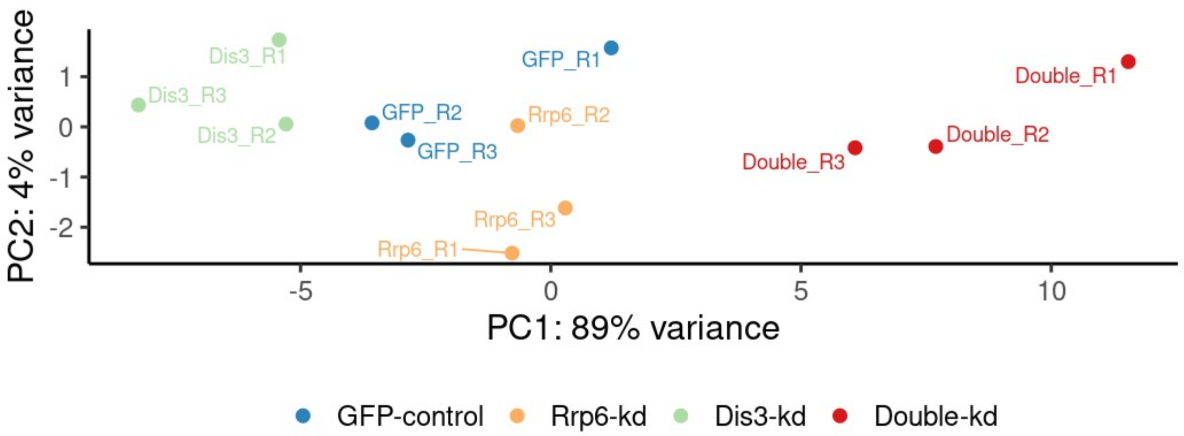


**Supplementary Figure S7. PCA analysis of ATAC-seq data in exosome depleted cells.** Principal component 1 (x-axis) separates the different RNA exosome depletion conditions into two groups. RRP6 and DIS3 are more similar to GFP control condition while the Doublekd is clearly differentiated.

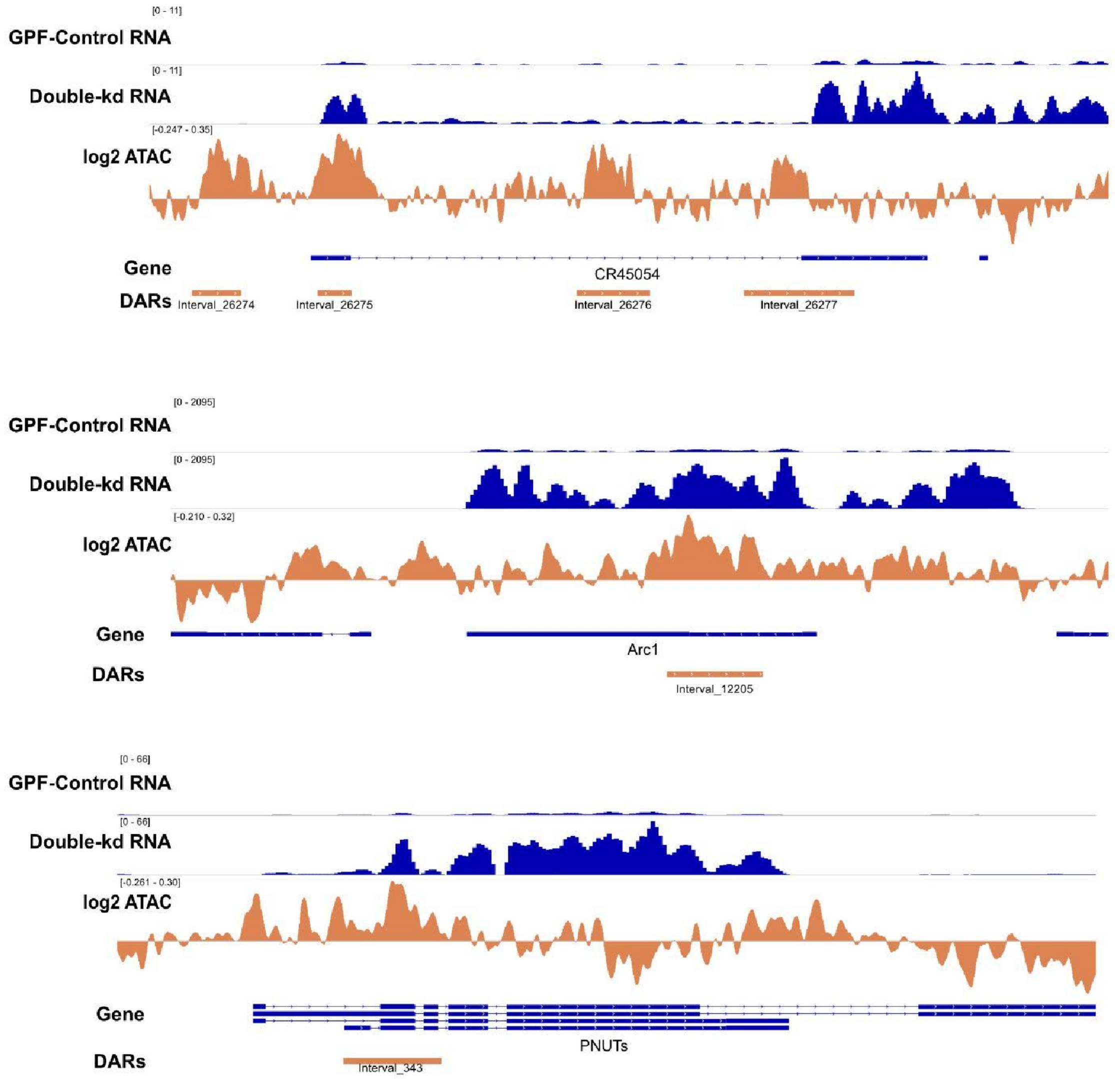


**Supplementary Figure S8. Three examples of DEG_DARs visualized in IGV.** Browser track visualization of RNA-seq signal in control-GFP and Double-kd (blue) and ATAC-seq change in Double-kd (log2 fold change Double-kd *vs*. GFP-control, orange). The signal was normalized by library sequencing depth (CPM).

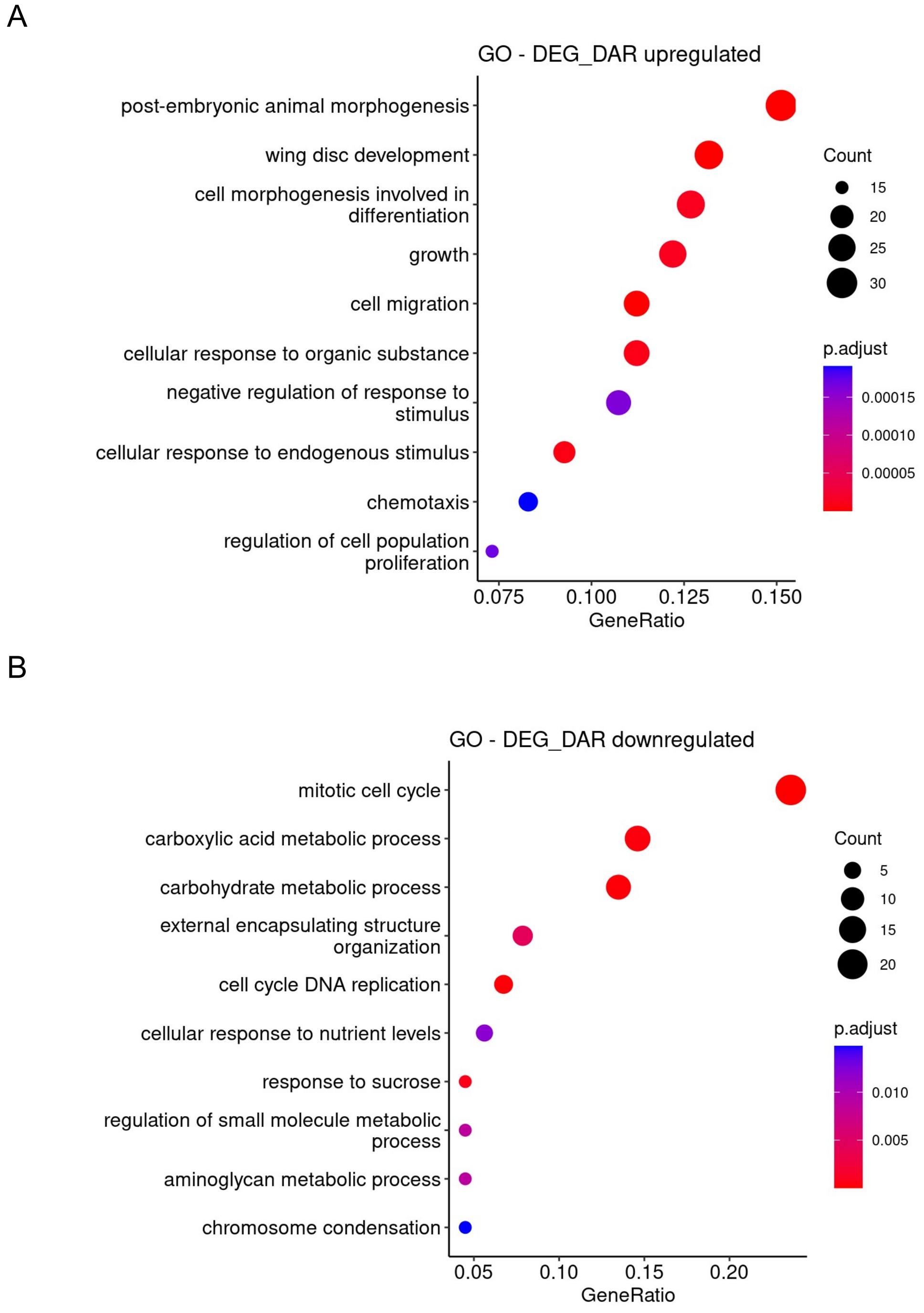


**Supplementary figure S9. GO of DEG_DARs increased and decreased in Double-kd.** In each case, the plot shown the top ten most significant categories in the y-axis. Statistical significance is color coded. The x-axis shows the relative amount of DE caRNA genes in each category. Dot size indicates the number of genes in each GO term. **(A)** and **(B)** show the biological processes associated to DEG_DARs that show increased or decreased levels in Double-kd compared to GFP-control, respectively.

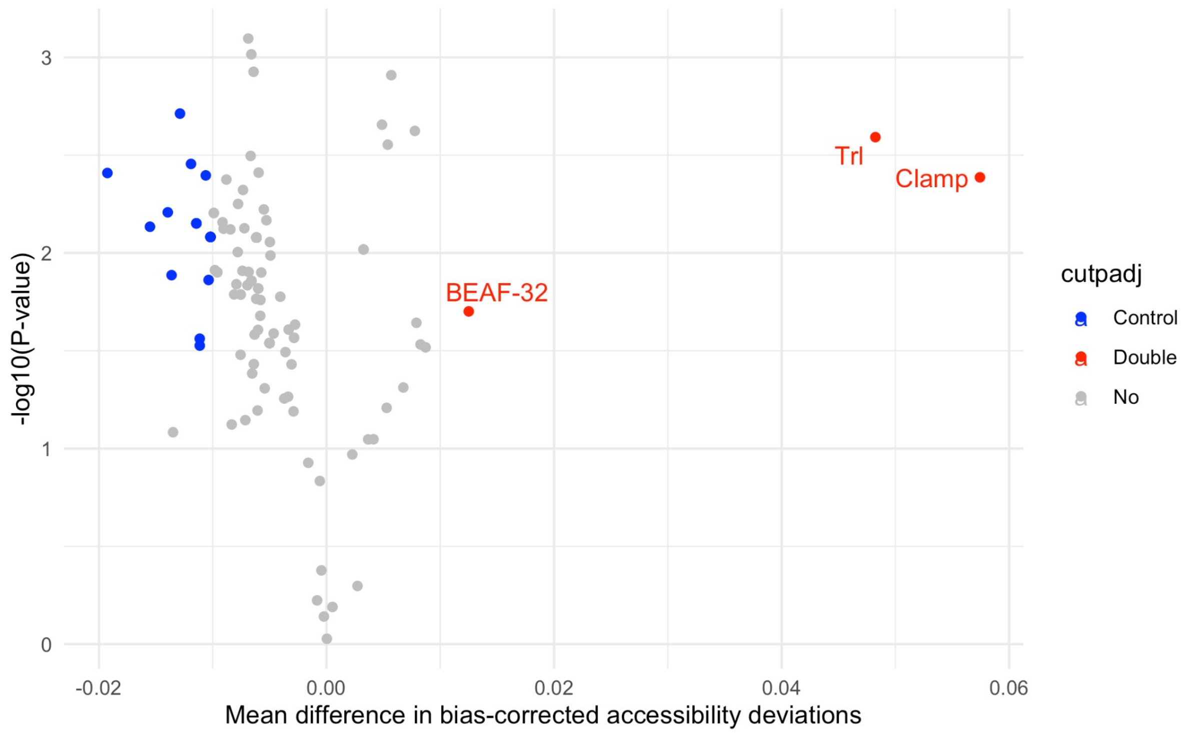


**Supplementary figure S10. Motifs enriched in DARs in Double-kd using ChromVar**. The plot shows accessibility deviations of DNA binding factors in Double-kd differential accessible regions. chromVar was used to compute accessibility deviations between groups (GFP-control and Double-kd, x-axis). Significance of the change is shown in y-axis. Blue colored dots correspond to DNA binding factors more prevalent in GFP-control condition, while red dots are binding factors enriched in double-kd. The accessibility deviations were normalized by a set of background peak sets matched for GC and average accessibility. All available *D. melanogaster* binding motifs were retrieved from JASPAR-CORE database and used for this analysis.

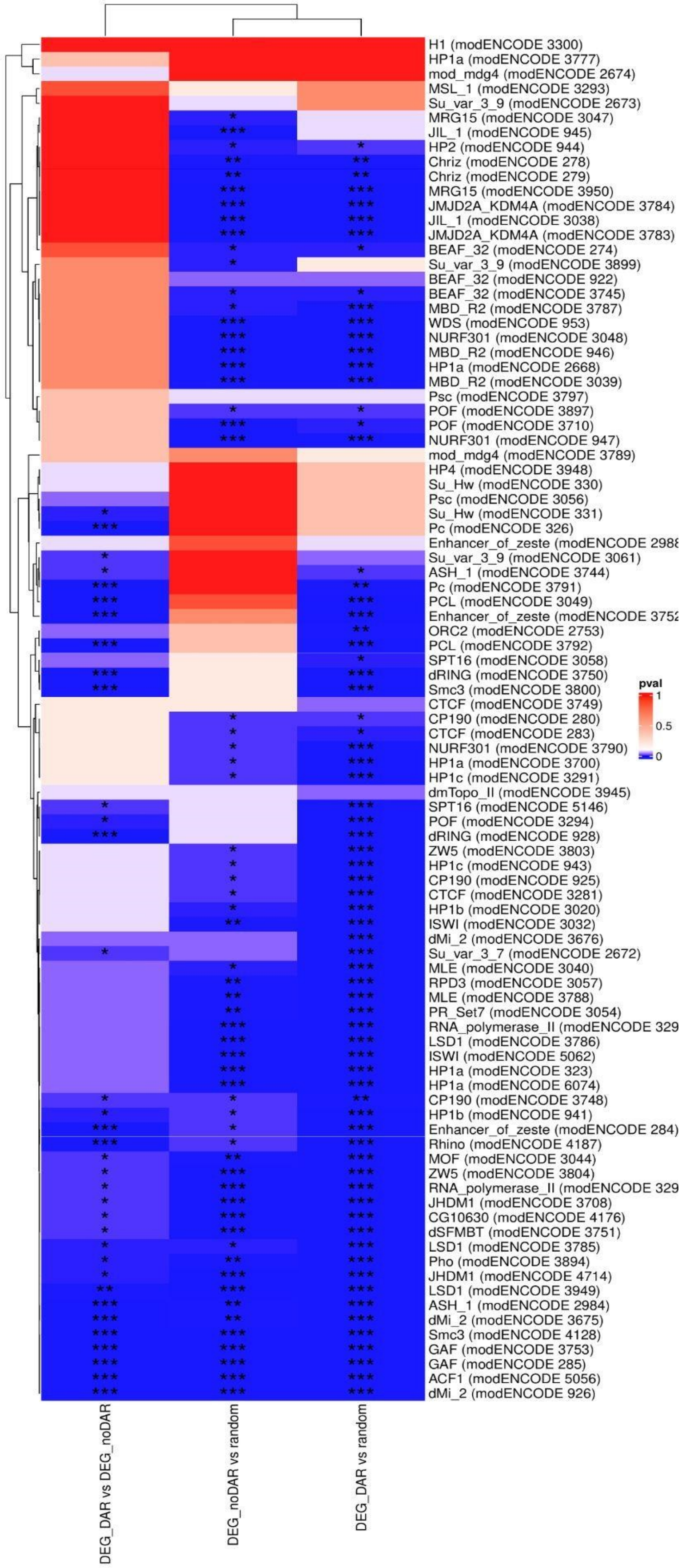


**Supplementary Figure S11. Analysis of modENCODE ChIP-on-chip data for 40 chromatin factors**. Heatmap showing the p-values of the statistical comparisons for modENCODE public data sets. The p-values have been calculated with the Kolmogorov-Smirnov test in a 1kb window around the TSS (+/- 500bp). Each row corresponds to a different modENCODE data set. The three columns correspond to the different comparisons analyzed for the three different gene sets (DEG_DAR vs Random, DEG_noDAR vs Random, DEG_DAR vs DEG_noDAR). Statistical significance is color coded, where dark blue corresponds to smaller p-values and higher pvalues are denote by red color (see color bar to the right of the heatmap).

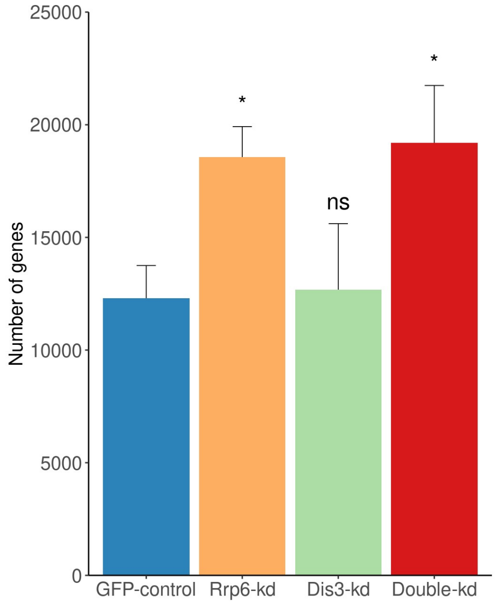


**Supplementary Figure S12. Number of detected rep-caRNAs in different experimental conditions**. The histogram shows the number of repetitive caRNAs detected in the different conditions, as indicated. The bars shows the average number of detected rep-caRNAs in the replicate experiments (n = 3). The error bars denote standard deviations. GFP has been used as reference in the two-sided Wilcox test for statistical significance. * indicates p<0.05. ns, not significant.

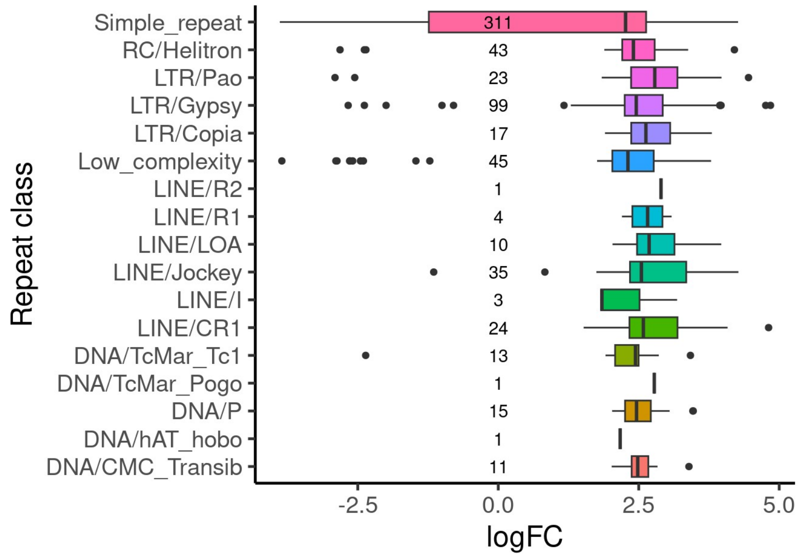


**Supplementary Figure S13. Classification of rep-caRNAs differentially expressed in exosome depleted cells**. The plot shows RNA abundance changes in Double-kd versus control GFP cells. The x-axis shows the log2FoldChange for each repeat class. Black line inside the box denotes the median for each repetitive class. Numbers next to the boxes indicate the number of DE rep-caRNAs differentially expressed in each repetitive class.
